# Sharks and rays have the oldest vertebrate sex chromosome with unique sex determination mechanisms

**DOI:** 10.1101/2025.03.13.642935

**Authors:** Taiki Niwa, Yoshinobu Uno, Yuta Ohishi, Mitsutaka Kadota, Naotaka Aburatani, Itsuki Kiyatake, Daiki Katooka, Michikazu Yorozu, Nobutaka Tsuzuki, Atsushi Toyoda, Wataru Takagi, Masaru Nakamura, Shigehiro Kuraku

## Abstract

Sex determination has been investigated across vertebrate lineages to reveal stepwise evolution of sex chromosomes and diversity of responsible molecular mechanisms. However, these studies hardly encompass cartilaginous fishes deeply isolated from the rest of vertebrates, which hinders the comprehensive view of vertebrate sex determination. Here, we produced chromosome-scale genome assemblies of egg-laying shark species and comparatively investigated genome sequences and transcriptome profiles across diverse cartilaginous fishes. Sex chromosome identification, supported by cytogenetic experiments, elucidated the homology of X chromosomes between sharks and rays as well as an extensively degenerating Y chromosome harboring no male-specific genes. These sex chromosomes scarcely included orthologs of previously documented sex determining genes. Transcriptomic analyses combined with histology of embryonic gonads revealed female-biased expressions of X-linked genes including those implicated in TGF-β and IGF signaling pathways, which are attributed to their incomplete dosage compensation. Our findings indicate that sharks and rays share the oldest vertebrate sex chromosomes that originated around 300 million years ago and the dosage-dependent sex determination mechanism comprised of distinct molecules from other vertebrates. This study highlights the antiquity of sex chromosomes and uniqueness of sex determination mechanisms in sharks and rays, which advance our understanding on evolutionary plasticity of vertebrate sex determination.

## Introduction

Sex chromosomes are prevalent among diverse mechanisms of sex determination, while they have been recurrently acquired at different timepoints in animal evolution^1^ (Fig. 1). In vertebrates, mammals and birds possess sex chromosomes with old origins, as evidenced by their genetic contents being widely shared within each of these taxa, which diverged over 100 million years ago^2–4^. These old origins are reflected in different sizes in their sex chromosome pairs (heteromorphy), characterized by massive gene loss on one chromosome in those pairs, owing to suppressed homologous recombination^2,5^. Mammalian X chromosomes further developed a compensation mechanism for their halved gene dosage, achieved by transcriptional inactivation of the one X chromosome in females^5^. On the other hand, teleost fishes^6,7^, frogs^8^, and geckos^9^ have homomorphic sex chromosomes with younger origins around several million years old. These observations were integrated into the concept of sex chromosome evolution involving cycles with different turning points^10–13^. A newly acquired sex chromosome pair is gradually differentiated by recombination suppression, leading to compensation of halved gene dosage. At some points, another chromosome overrides the sex determining function, resulting in its turnover. However, this model cannot encompass some observations. Birds and snakes exhibit incomplete dosage compensation on their heteromorphic ZW chromosomes^14,15^, prompting discussion on the necessity of dosage compensation^16,17^. Other recent findings in ratite birds^18^, sturgeons^19^, and scallops^20^ uncovered long-lasting homomorphy of their sex chromosomes. These exceptions provided important insights into our general understanding on sex chromosome evolution, which can be further refined by exploring the considerable number of unexamined taxa.

**Fig. 1:**
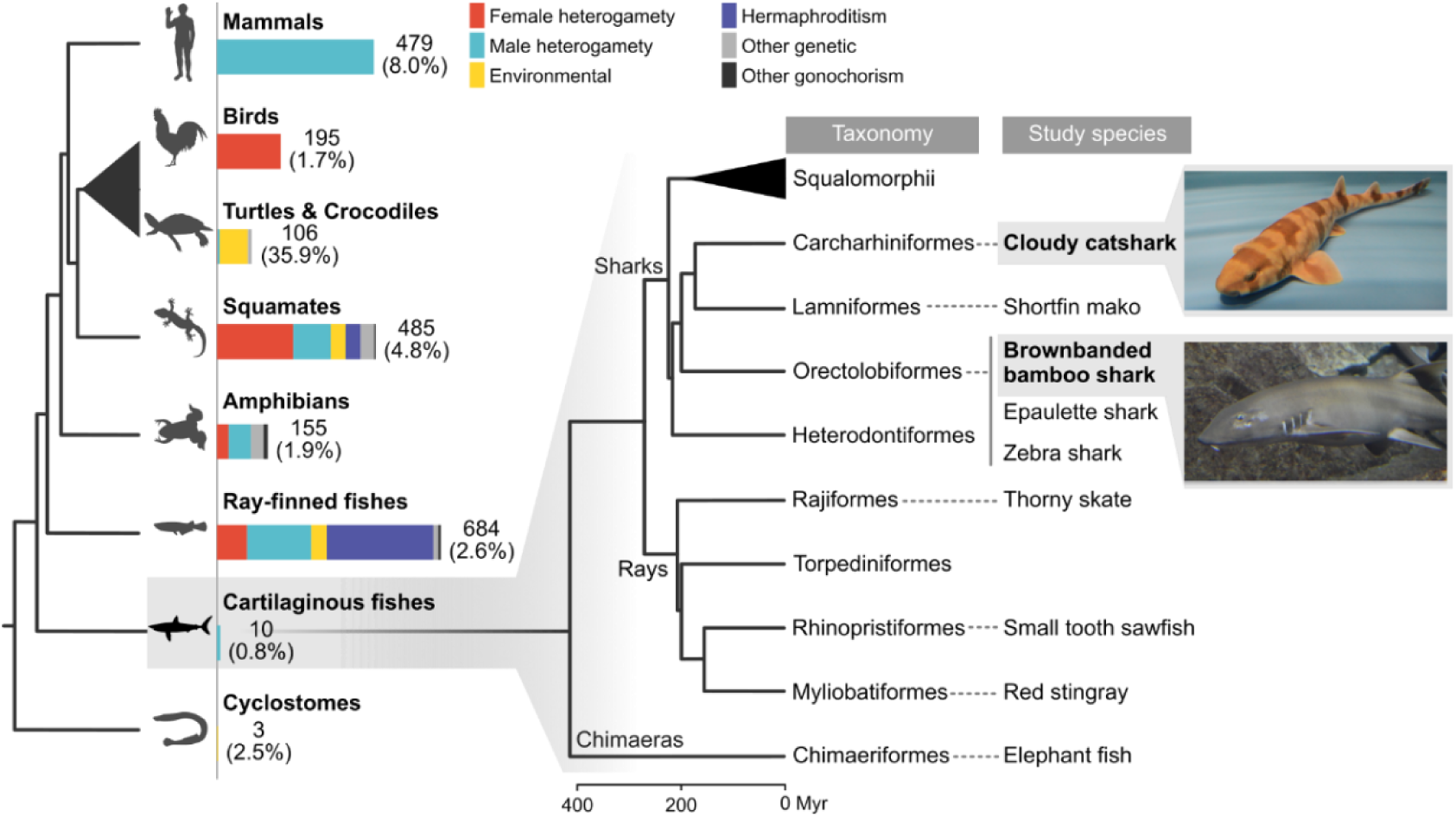
Our study systems to supplement the knowledge of vertebrate sex determination. (Left) Numbers and breakdowns of the records of sex determination mechanism for vertebrate classes^21^ with its proportion in the total number of species in brackets (adopted from IUCN Red List; https://www.iucnredlist.org/statistics). (Right) Order-level phylogeny of cartilaginous fishes and our study species. For simplicity, multiple shark orders are collapsed into the superorder Squalomorphii. Divergence times were adopted from TimeTree (https://timetree.org/). Silhouettes were obtained from PhyloPic (http://phylopic.org). Myr: million years.

One of such taxa is cartilaginous fishes (Chondrichthyes), including sharks, rays and chimaeras, which diverged from bony vertebrates (Osteichthyes) around 450 million years ago (mya) (Fig. 1). While knowledge of sex chromosomes about bony vertebrates is gradually accumulating^21^, those in cartilaginous fishes were poorly characterized. This limitation is mainly attributable to their enlarged genome size in addition to occasional sample availability^22^. These difficulties are reflected in scarce cytogenetic reports on cartilaginous fishes^21,23^. On the other hand, advancements in sequencing technologies have significantly reduced the barriers to assembling genomes on the chromosome level^24,25^. Notably, recent studies revealed the presence of male heterogametic sex chromosomes in three shark species by assembling their genomes^26,27^, which was consistent with earlier cytogenetic observations^23^. These genomic studies pointed out small pseudoautosomal regions (PARs) or a highly degenerated Y chromosome, which implies their ancient origins. Research on the white-spotted bamboo shark further demonstrated the absence of global dosage compensation on its X chromosome^27^. Although these pioneering studies unveiled distinct feature of sex chromosome in cartilaginous fishes, they are limited to three species from the order Orectolobiformes, one of the 14 orders in cartilaginous fishes (Fig. 1). Consequently, the entire evolutionary history of sex chromosomes in cartilaginous fishes remains unsolved.

Besides these evolutionary aspects, sex determination have been investigated by developmental biologists. In many vertebrates, a master sex determining gene on a sex chromosome starts to canalize embryonic gonads into either a testis or an ovary, by its sexually biased transcription^28^. While mammals and birds only employ *Sry* (*Sox3* homolog) and *Dmrt1*, teleost fishes adopt sex determining genes with various functions including transcriptional factors (*Sox3* and *Dmrt1* homologs), TGF-β ligands and their receptors (*Amh*, *Gsdf*, *Gdf6*, and *Amhr2* homologs)^7,28^. These differences in availability of potential sex determining genes seem to be associated with the different turning points of their sex chromosome evolution; frequent turnovers in teleosts and rare turnover in mammals and birds. In contrast to the arising evolutionary studies^26,27^, the molecular basis of sex determination remains to be elucidated in cartilaginous fishes, because of the limited availability of embryos owing to their low fecundity and predominant viviparity^29,30^.

To untangle the evolutionary history and molecular basis of sex determination in cartilaginous fishes, we resorted to their emerging genomic resources as well as embryonic materials from egg-laying sharks with high fecundity. Our genomic comparison encompassing diverse cartilaginous fishes revealed the ancient origin of their X chromosomes around 300 mya, representing the oldest origin among vertebrate sex chromosomes, in agreement with the extensive degeneration of their Y chromosomes. Transcriptomic analyses demonstrated the absence of global dosage compensation at embryonic gonads as well as livers, which possibly underlies their sex determination mechanism. Our findings highlight the antiquity of elasmobranch sex chromosome and advance our understanding on the evolutionary plasticity of vertebrate sex determination.

## Results

### Chromosome-scale genome assembly of two shark species

We focused on two shark species, the brownbanded bamboo shark (*Chiloscyllium punctatum*) and the cloudy catshark (*Scyliorhinus torazame*) (see images in Fig. 1) that can be bred in captivity, allowing the constant access to fertilized eggs. Previously, their genome sequencing was performed only on non-chromosomal levels^31^. In the present study, we produced chromosome-scale genome assemblies of these species, incorporating long and accurate genomic reads and Hi-C scaffolding. Over 90% of our assemblies in length were composed of chromosome-scale scaffolds that were larger than 10 Mb and enriched with intra-chromosomal chromatin contacts (Fig. 2a). The numbers of the chromosome-scale scaffolds were 53 and 32 in the bamboo shark and the cloudy catshark, respectively, which matched the numbers of the chromosomes in their karyotypes (2n = 106 and 2n = 64, respectively; refs 23,32) (Fig. 2a). The assembly sizes of these two genomes were comparable to the nuclear DNA content estimated by flow cytometry: 4.73 Gb for the bamboo shark and 6.73 Gb for the cloudy catshark^31^ (Fig. 2b). These assemblies were assessed by the retrieval of single-copy orthologs conserved among vertebrates, which marked the completeness of more than 90% in both species (Fig. 2c). On these genome assemblies, we inferred protein-coding gene models, which contained over 90% of the conserved ortholog set (Fig. 2d). Over 98% of the inferred genes were located on the chromosome-scale scaffolds. These metrics indicate that the genome assemblies obtained in this study are chromosome-scale, reflecting the actual genomic organization.

**Fig. 2:**
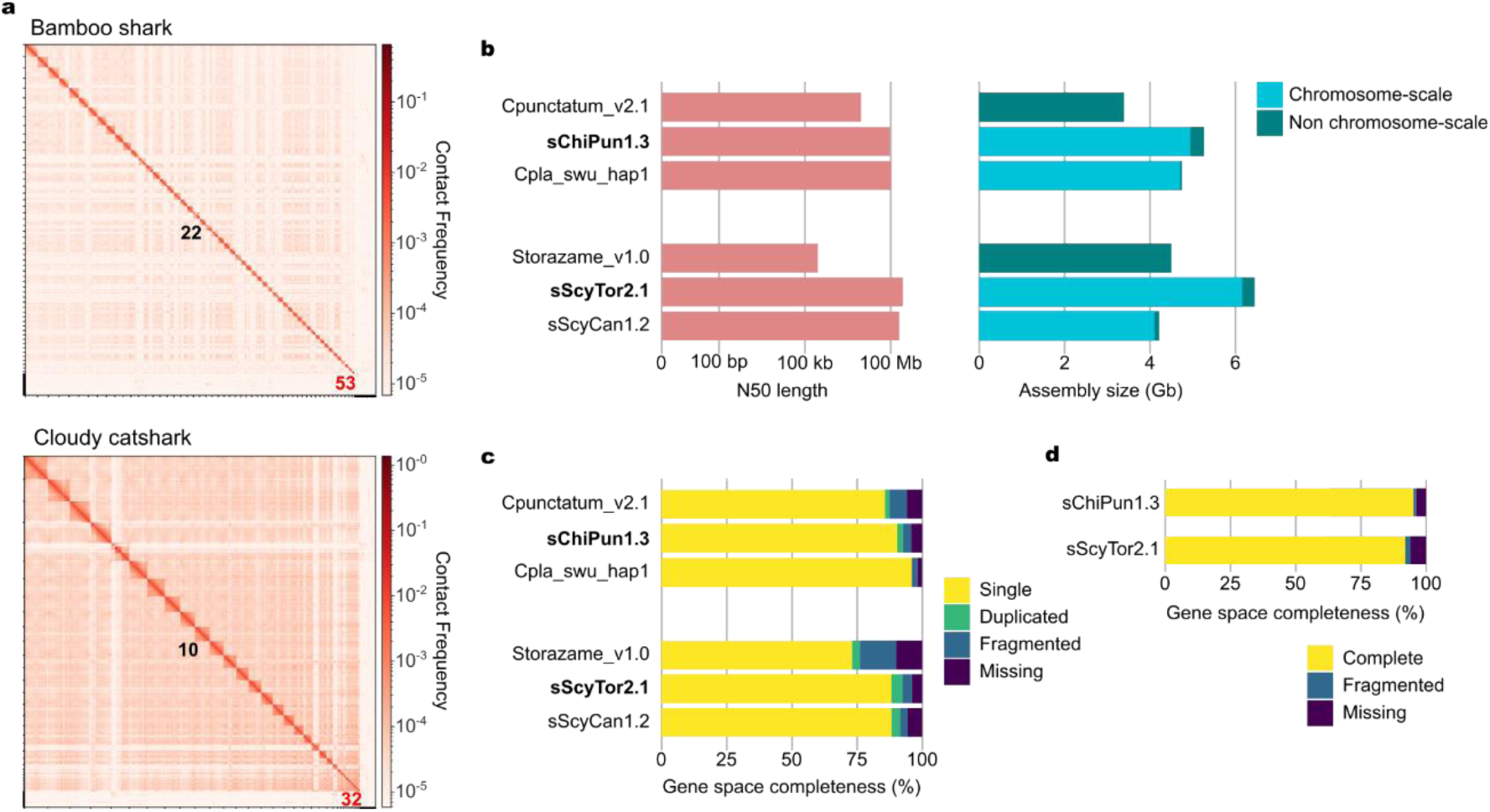
Assessment of our chromosome-scale genome assemblies of two shark species. **a**, Genome-wide contact map based on Hi-C data. Scaffolds are ordered by length. Numbers in black and red indicate L50 scaffolds and the smallest chromosome-scale scaffolds, respectively. **b**. **c**, N50 length, assembly size, occupancy of chromosome-scale scaffolds (**b**) and gene space completeness (**c**) of the previous assembly, new assembly (bold) and the chromosome-scale assembly from closely-related species (Cpla_swu_hap1: *Chiloscyllium plagiosum*, sScyCan1.2: *Scyliorhinus canicula*). **d**, gene space completeness of our gene models. See Methods for the details of gene space completeness analysis.

### Homology between shark and ray X chromosomes

To identify X chromosome sequences, we compared the genomic short-read mapping data between sexes utilizing original and public genome assemblies. These datasets were derived from both sexes of four shark, three ray and one chimaera species that encompasses the entire diversity of cartilaginous fishes^25,33–36^. In a differentiated sex chromosome pair, the male X chromosome is expected to show a halved sequencing depth and low heterozygous site frequency (heterozygosity), compared with autosomes, because of its male-specific hemizygous pattern. In our datasets, six species including sharks and rays possessed single chromosomes that showed the male-female differences featuring X chromosomes (Fig. 3a and Extended Data Fig. 1). X chromosomes originally registered in the epaulette shark and the small tooth sawfish genomes were confirmed again in our analysis, while that of the thorny skate was not (Extended Data Fig. 1). In the thorny skate, we identified the original X chromosome as an autosome, and instead, original chromosome 46 as an X chromosome (Extended Data Fig. 1).

**Fig. 3:**
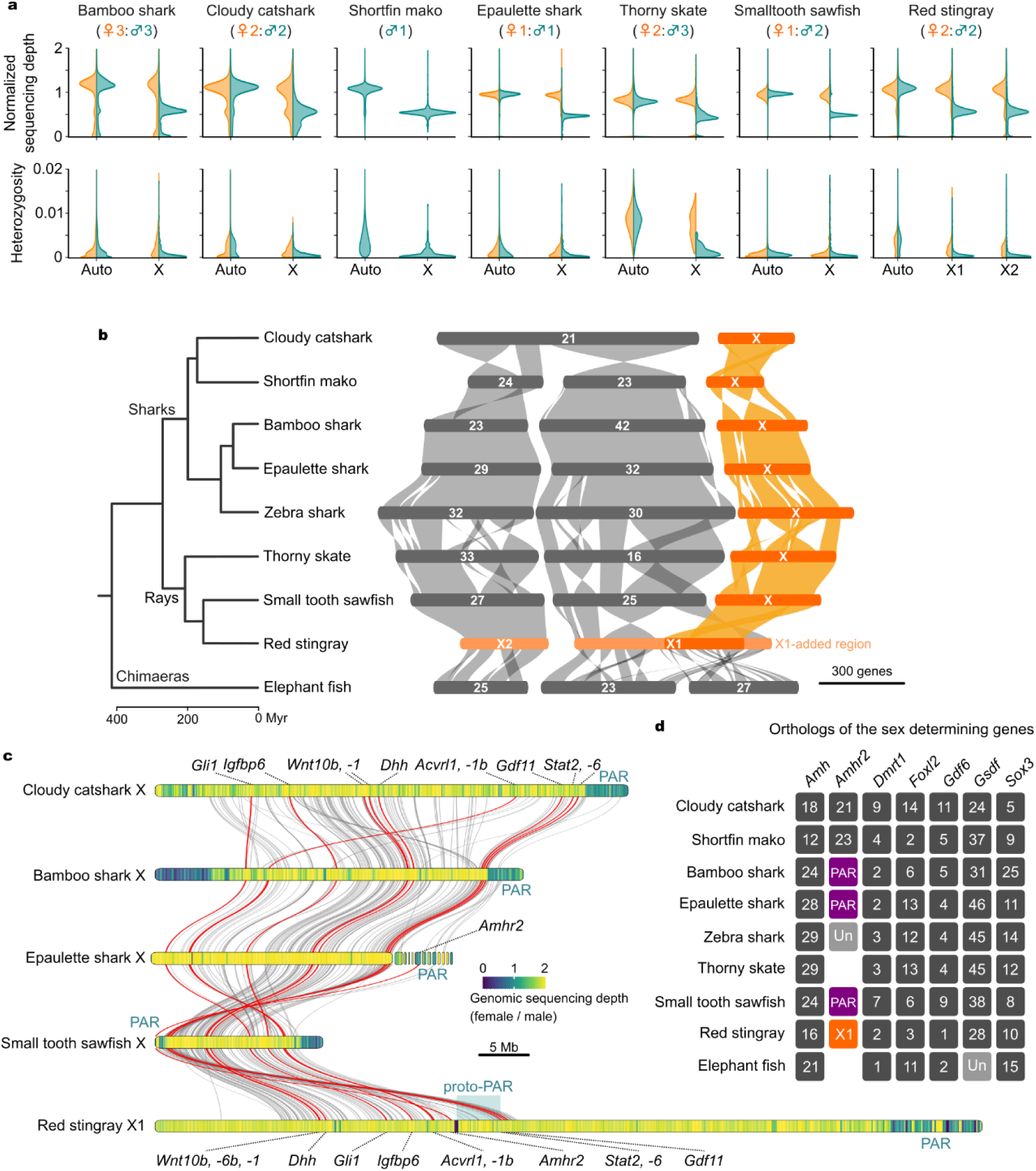
Identification of X chromosomes and their homology. **a**, Distribution of normalized sequencing depth and heterozygous site frequency (heterozygosity) of chromosomes (‘X’ and ‘Auto’). These two metrics were computed for non-overlapping windows of 10 kb and 100 kb, respectively, from whole genome resequencing of male and female individuals. Sequencing depth was normalized by its whole-genome average. Data for shortfin mako (*Isurus oxyrinchus*), epaulette shark (*Hemiscyllium ocellatum*), smalltooth sawfish (*Pristis pectinata*), and thorny skate (*Amblyraja radiata*) are derived from public repositories. **b**, Macrosynteny-based homology of X chromosomes. Divergence times were adopted from exiting literature (https://timetree.org/; ref. 37). **c**, Microsynteny-based comparison among X and X1 chromosomes in selected species. Red lines indicate the orthology for the twelve genes annotated at the bottom. The color spectrum (navy to yellow) shows female-to-male ratios of genome sequencing depth in 100-kb non-overlapping windows. **d**, Genomic location of elasmobranch orthologs of master sex determining genes documented in other animals. Numbers and ‘X’ indicate chromosome identities. ‘Un’ and blank stand for unplaced scaffold and no ortholog detected, respectively. See Extended Data Fig. 4 for the details of orthology assessment.

Our comparison revealed two distinct chromosomes of red stingray (*Hemitrygon akajei*, designated as X1 and X2) with the same features as those in the other species. In the currently available genome assembly of the elephant fish (*Callorhinchus milii*), a chimaera, none of the chromosome-scale scaffolds showed the hemizygous pattern (Extended Data Fig. 1), showing the absence of the heteromorphic sex chromosome in this lineage.

Utilizing the identified X chromosome sequences and that of the zebra shark (*Stegostoma tigrinum*)^26^, we examined homology among these chromosomes to trace their evolutionary history. All the X chromosomes of sharks and rays, including a part of X1 chromosome in the red stingray, harbored the common gene repertoires with several rearrangements (Fig. 3b). This suggests that the origin of these X chromosomes and their sex determining function can be traced back to the common ancestor of elasmobranchs (sharks and rays), at least.

The red stingray possessed an additional X chromosome (X2) as well as a large region of the X1 chromosome which did not share gene repertoires with the X chromosome of other elasmobranch species (X1-added region, Fig 3b). Each of these regions was homologous to an autosome in the other elasmobranch species (Fig. 3b). This X1/X2 chromosomal organization was reported in an earlier cytogenetic study in the genus *Potamotrygon*^38,39^ and later in the genome assembly of the Atlantic stingray (*Hypanus sabinus*, GCF_030144855.1). These species are categorized into the same order Myliobatiformes as the red stingray, supporting our finding in the red stingray. Although assignments of sex chromosome in the Atlantic stingray were not clearly evidenced, the X1 and X2 chromosomes showed the conserved synteny between the red stingray and the Atlantic stingray (Extended Data Fig. 2). Y-linked genes in the Atlantic stingray retained their homologs (gametologs) on both X1 and X2 chromosomes, indicating that this Y chromosome have its origin at both of the X1 and X2 chromosomes. These observations imply that the myliobatiform X1 and X2 chromosomes were estimated to have been established by two chromosomal rearrangements: a fusion between proto-X and autosome resulting in X1, and a fusion between proto-Y and autosome resulting in X2.

### Compositions of the X chromosomes

Heteromorphic sex chromosomes, such as mammalian XY chromosomes, maintain recombination only in PARs, resulting in their high sequence similarity between sex chromosome pairs. We found the small region featuring PAR in the five species of sharks and rays, as the X chromosome ends sequenced at comparable depths between sexes in whole genome resequencing (Fig. 3c and Extended Data Fig. 3). These PARs shared the subset of gene repertoires including Hox C cluster, *Hnrnpa1*, *Cnpy2*, *Cldn12*, and *Slc39a5* in current gene models. These regions tend to show relatively high GC-content compared with the remainder of their X chromosomes, which is consistent with general features of PARs originating from their high recombination frequency^40–43^ (Extended Data Fig. 3). Considering these common features, single ends of the X chromosomes in the bamboo shark and small tooth sawfish were regarded as PARs in the following analyses. The sizes of the PARs were much smaller than those of entire X chromosomes, indicating extensive differentiation of these sex chromosomes (Fig. 3c). Unlike the other species, the PAR in the red stingray X1 chromosome was located in the X1-added region, which was homologous to the autosomes in the other species (Fig. 3b, c, and Extended Data Fig. 3). Although the region homologous to the PARs in the other species exhibited the relatively high GC-content, this region showed the female-biased sequencing depth in the red stingray (hereafter referred to as proto-PAR, Fig. 3b).

By means of this high-completeness dataset, we searched for cartilaginous fish orthologs of the seven master sex determining genes that were documented in independent animal lineages: *Amh* (pike and pejerrey), *Amhr2* (fugu, cichlid, and catfish), *Dmrt1* (Japanese medaka, birds, and western clawed frog), *Foxl2* (Chinese crocodile lizard and scallops), *Gdf6* (Mexican tetra and turquoise killifish), *Gsdf* (fugu and medaka *Oryzias luzonensis*), and *Sox3* (placental mammals and Indian medaka)^7,20,44^. Our molecular phylogenetic assessment of putative cartilaginous fish homologs validated their orthology, but these orthologs were not identified on the elasmobranch X chromosomes except for *Amhr2* (Fig. 3c and Extended Data Fig. 4a-c).

The *Amhr2* orthologs were located on the PARs of the epaulette shark and the small tooth sawfish, the proto-PAR of the red stingray (Fig. 3c, d), and autosomes of the cloudy catshark and the shortfin mako (Fig. 3d). The bamboo shark *Amhr2* ortholog was only included in raw genomic reads and the transcriptome assembly reported previously^31^. Sequencing depths of the coding region of this gene exhibited no marked difference either from autosomal regions or between sexes (1.05 for males versus 0.934 for females in normalized depth of whole genome resequencing). Taking advantage of our Hi-C data, we confirmed that the raw HiFi genomic read including this gene showed the most frequent contact with the X chromosome (Extended Data Fig. 4d), indicating that this gene was located on the bamboo shark PAR. These results imply that *Amhr2* in sharks and rays is not specifically localizaed on either X or Y chromosome.

Sex determination cascades, triggered by master sex determining genes, involve numerous intercellular signalling pathways, including TGF-β, Wnt, Hedgehog, IGF, and JAK/STAT signaling pathways^45–54^, leading to sexual differentiation of gonads. We further searched for the X-linked genes that potentially have the function in these intercellular signaling pathways and identified eleven genes commonly harbored in X chromosomes of five species analyzed here: TGF-β signaling pathway (*Acvr1b*, *Acvrl1*, and *Gdf11*), Hedgehog signaling pathway (*Dhh* and *Gli1*), Wnt signaling pathway (*Wnt1*, *-6b*, and *-10b*), IGF signaling pathway (*Igfbp6*), and JAK/STAT signaling pathway (*Stat2* and *Stat6*) (Fig. 3c and Extended Data Fig. 5).

### Y chromosome identification

We revealed the extensive differentiation of the elasmobranch sex chromosome. To uncover male-specific elements on the Y chromosomes, we aimed to identify the Y chromosome sequences from the bamboo shark genome. We dissected Y chromosomes on chromosome spreads using fine glass needles, sequenced them and mapped onto the reference genome. The validity of the chromosome dissection was examined by fluorescence *in situ* hybridization (FISH) on chromosome spreads (Fig. 4a). The genomic sequences from the chromosome dissection were most enriched in a single scaffold (scaffold 59) (Fig. 4a). This scaffold was sequenced specifically in male individuals in the whole genome resequencing and exhibited relatively high chromatin contact frequency with the PAR identified as a tip of the X chromosome (Fig. 4b and Extended Data Fig. 6a). These results suggest that this male-specific scaffold and the PAR represent the bamboo shark Y chromosome and were concatenated for following analyses. For the epaulette shark, one chromosome-scale and two unplaced scaffolds were originally registered as the Y chromosome in its genome assembly, but with no detailed evidence. We checked their depth in whole genome resequencing, and one chromosome-scale scaffold was confirmed as the male-specific scaffold, and the other two scaffolds were confirmed as fragments of the PAR (Fig. 4b).

**Fig. 4:**
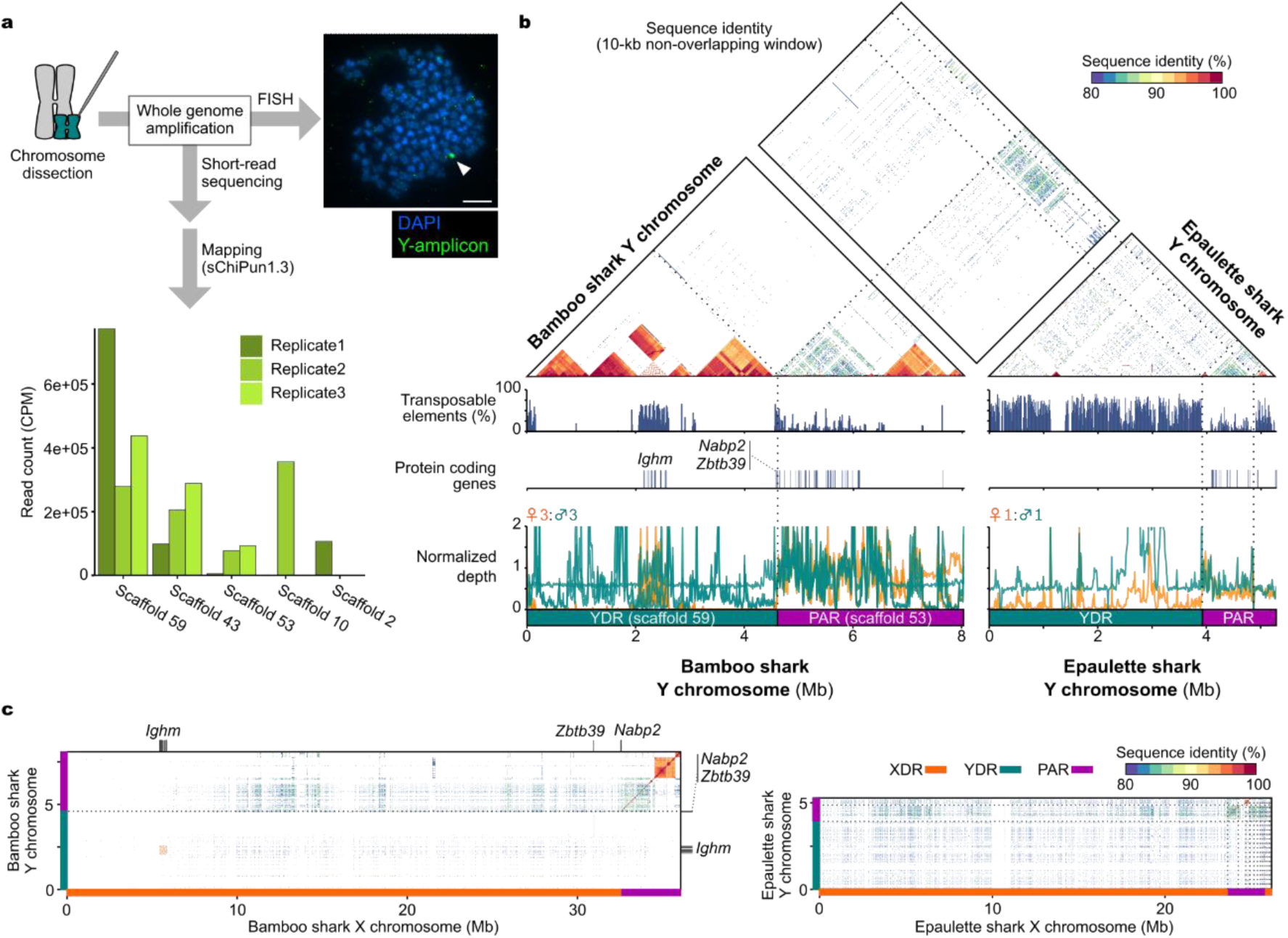
Identification of shark Y chromosomes and their degeneration. **a**, Identification of the bamboo shark Y chromosome. Metaphase spreads were prepared from male bamboo shark fibroblasts and hybridized with DNA probes derived from the Y chromosome dissection. Arrowhead, Y chromosome. Scale bar, 10 μm. The bar graph indicates enrichment of sequencing reads derived from Y chromosome dissection. Top five scaffolds are shown. **b**, Sequence features of Y chromosomes of two shark species. Normalized depth of whole genome resequencing and interspersed repeat coverage were quantified for10 kb non-overlapping windows. Sequence similarities were computed between all pairs of 10 kb non-overlapping windows. **c**, Sequence similarity between X and Y chromosomes in two shark species. Locations of bamboo shark X-Y gametologs were shown as black lines on the edge of the plot. Dotted lines in **b** and **c** indicate the borders of concatenated sequences. See Extended Data Fig. 6c for the equivalent plot for the white-spotted bamboo shark.

We analyzed the sequence similarity between X and Y chromosomes in the two shark species to assess the degree of sex chromosome differentiation. In agreement with the limited PAR extent on the X chromosomes, the regions specific to either the X or Y chromosomes (hereafter referred to as X/Y-differentiated region, abbreviated in XDR/YDR) no longer shared common sequences along their entire extent (Fig. 4c).

According to our gene model, the YDR in the bamboo shark harbored nine protein-coding genes including a tandem cluster of *Ighm* (immunoglobulin heavy chain mu), *Zbtb39* (Zinc finger and BTB domain-containing protein 39), a partial sequence of *Nabp2/Ssb2* (Nucleic acid binding protein 2 / Single strand binding protein B1), and one small uncharacterized gene (Fig. 4b). All these genes had their gametologs on the XDR, which exhibited high similarity between the X and Y chromosomes (over 94% identity in aligned peptide sequences). Although the allelic correspondence of *Ighm* genes were not fully resolved, both X and Y clusters contained six *Ighm* genes. These observations indicate the recent divergence of these gametologs. The epaulette shark YDR was devoid of any protein coding genes according to the gene annotation provided by NCBI (Fig. 4b). Based on our repetitive sequence annotation, YDR of the two species were occupied by different types of repetitive sequences: satellite repeats in the bamboo shark (43-108 bp/unit) and transposable elements in the epaulette shark (Fig. 4b and Extended Data Fig. 6b). Interspecific sequence comparison revealed the absence of conserved sequences among the YDRs of three shark species, including the white-spotted bamboo shark, a more closely related species to the bamboo shark than the epaulette shark (Fig. 4b and Extended Data Fig. 6b). These results indicate that the Y chromosomes in these species lost a large fraction of their ancestral genomic contents and have independently accumulated different types of repetitive sequences.

### Dosage-dependent expression of X-linked genes

Sex chromosome degeneration is often accompanied by the acquisition of dosage compensation. We examined this on the X chromosomes of three elasmobranch species (the bamboo shark, the cloudy catshark and the red stingray) using RNA-seq data from the liver. The gene-by-gene ratio of the expression level between sexes was biased approximately 1.5 times towards females on their X chromosomes, in contrast to the sexually comparable expression of the autosomal genes (Fig. 5a and Extended Data Fig. 7a). This bias was also observed on both X1 and X2 chromosomes in the red stingray on the same level (Extended Data Fig. 7a). X-linked genes exhibited a lower expression level than autosomes only in males, implying the reduced expression of X-linked genes in males from their ancestral levels estimated from autosomal genes (Fig. 5b). The level of the expression bias between sexes was comparable to that of Z-linked genes in the chicken^55^, which lacks global dosage compensation, and this bias was absent on the compensated X-linked genes in the mouse^56^ (Extended Data Fig. 7a). The level of female-biased expression was constant along the position on their X chromosomes, except for their PARs (Extended Data Fig. 7b). Genes on the PARs, in which gene dosage is equivalent between sexes, tended to be expressed on the comparable level between sexes (Fig. 5a, Extended Data Fig. 7b). These trends were more apparently observed in other tissues; bamboo shark gonadal primordia and red stingray kidneys^57^ (Extended Data Fig. 7c). These observations suggest that sharks and rays lack dosage compensation on the chromosome level, and the expression level of X-linked genes is passively influenced by halved gene dosage in male, rather than actively regulated.

**Fig. 5:**
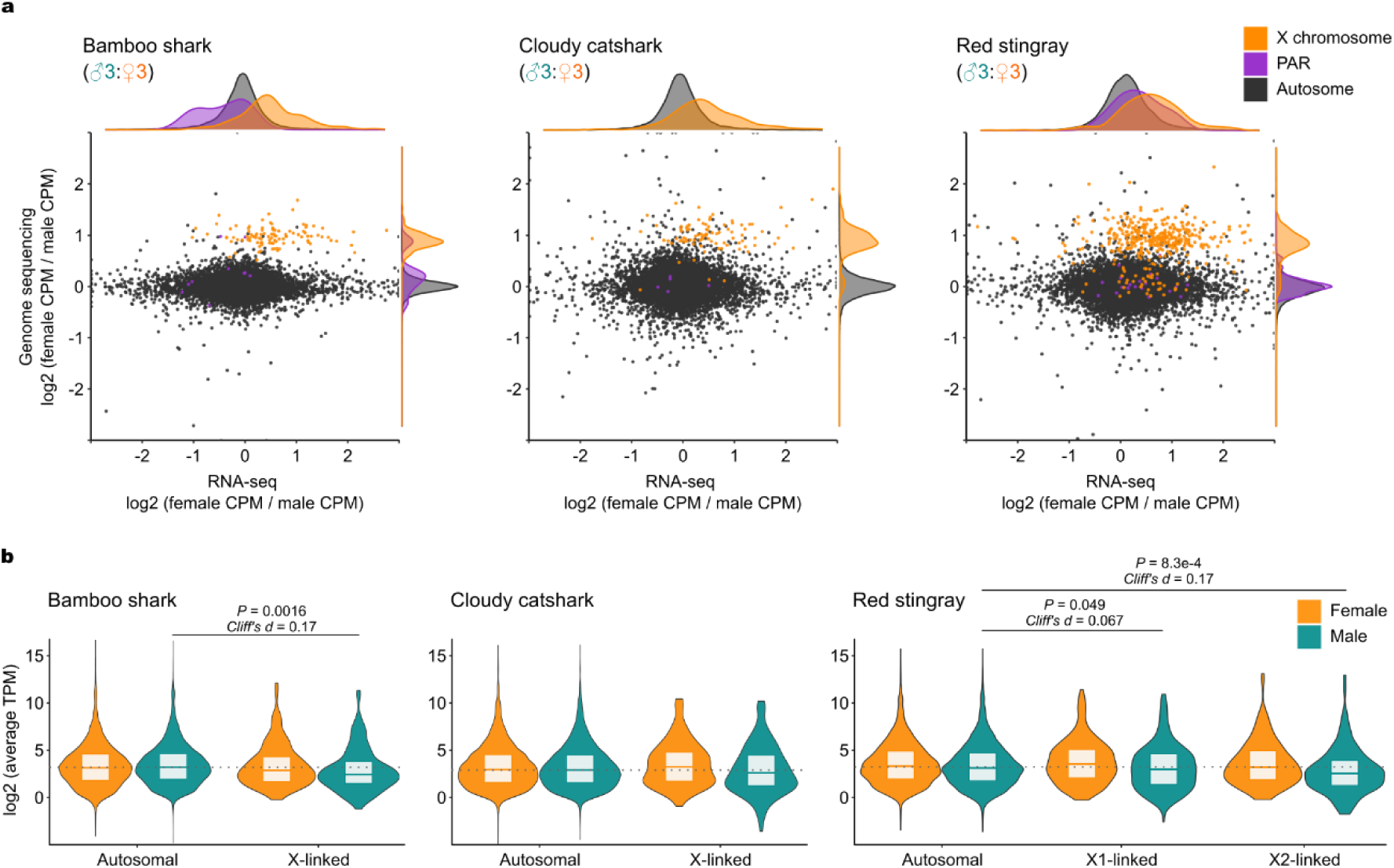
Dosage dependent expression of X-linked genes. **a**, Sexual biases in gene expression and gene dosage in the RNA-seq of livers from three elasmobranch species. Genes, represented by dots, were classified by different categories of their locations: autosome, XDR (X chromosome), and PAR. Each dot represents a gene. **b**, Distributions of the gene expression levels quantified from RNA-seq of livers from the three species. Values are compared by Wilcoxon rank sum test. Boxplot elements are defined as follows: center line, median; box limits, upper and lower quartiles. All values are averaged for individuals of each sex for each gene.

### Sex-biased gene expression in gonadal development

To untangle the molecular basis of the sex determination, we focused on sexual differences in the transcriptome of gonadal primordia. Previous studies revealed that the primordia of claspers, a pair of male copulatory organs, initially formed around the stage 30 bamboo shark embryo as a pair of thickenings on pelvic fin tips^58^.

Concomitantly, our histological observations on bamboo shark embryos revealed that gonadal primordia and reproductive tracts became sexually dimorphic around the stage 31; the right gonadal primordium extended, its border with the epigonal organ became prominent, and both sides of Müllerian ducts rapidly enlarged in a female-specific manner (Fig. 6a, b, and Extended Data Fig. 8). The sexual dimorphism in reproductive tracts was also evidenced in the cloudy catshark and the red stingray^59^. These observations implied that gonadal sex determination occurred at around stage 31, which instructed us to compare the transcriptomes at this stage.

**Fig. 6:**
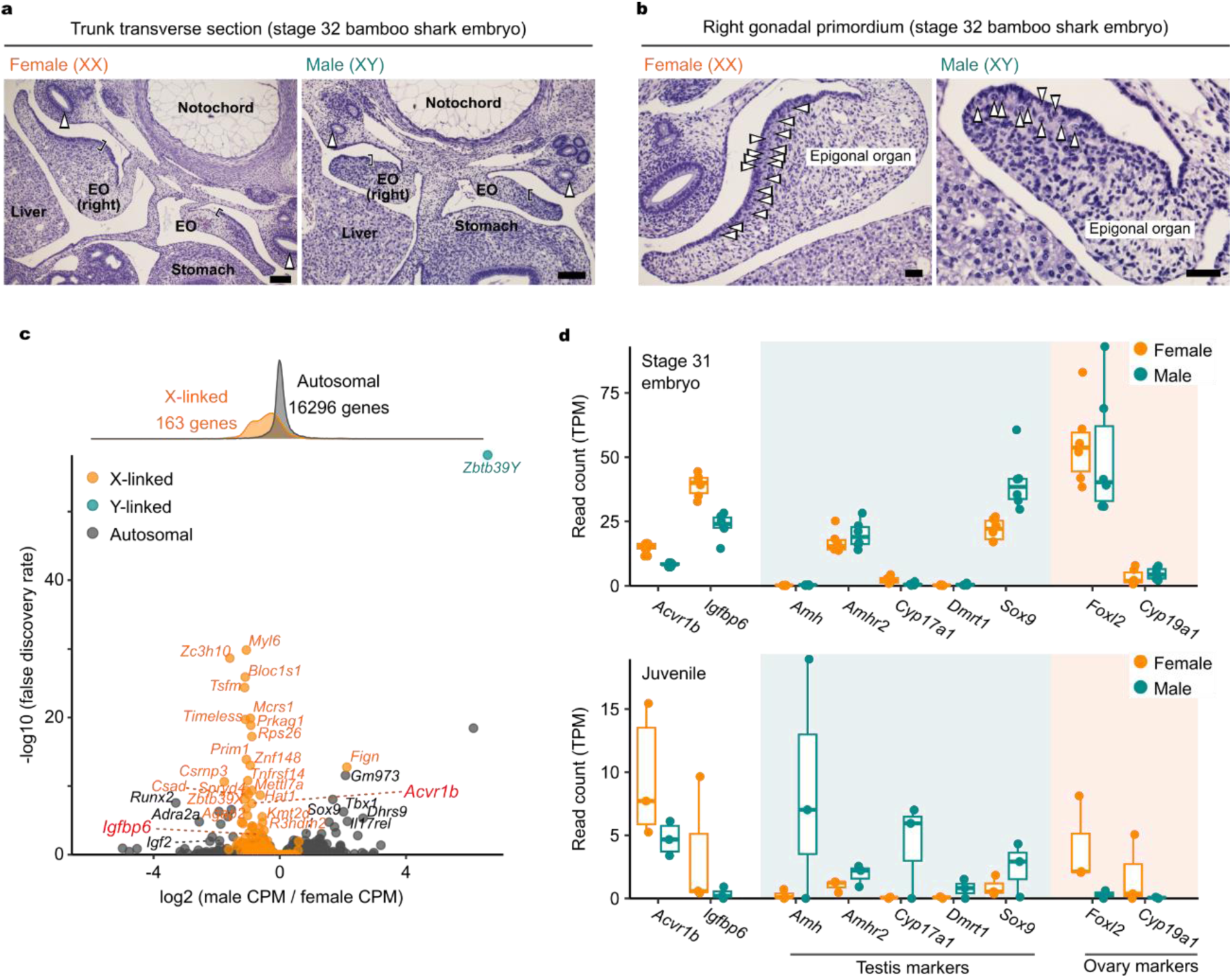
Sexual differences in morphology and gene expression of bamboo shark developing gonads. **a**, Transverse sections of bamboo shark trunks at the embryonic stage 32 stained with hematoxylin and eosin. **b**, Magnification from right gonadal primordia in **a**. EO: epigonal organ, arrowheads in **a**: Müllerian duct, brackets: gonad, arrowheads in **b**: germ cell. Scale bars, 100 μm (**a**), 40 μm (**b**). **c**, Differentially expressed genes between sexes in gonadal primordia of stage 31 bamboo shark embryos. See Methods for details of gene annotation. **d**, Expression levels of *Acvr1b* and *Igfbp6* as well as genes well-characterized for their sex-specific role. Data are derived from stage 31 embryos and juvenile individuals. Boxplot elements are defined as follows: center line, median; box limits, upper and lower quartiles; whiskers, maximum and minimum within 1.5×IQR from hinges.

We dissected gonad-epigonal organ complexes from bamboo shark individuals at embryonic stage 31 as well as juvenile stage, which were used for RNA-seq. These data included transcriptomic reads from germline-specific transcripts (e.g. *Ddx4* and *Dazl*), confirming the validity of the experiment (Extended Data Fig. 9a). The female-biased expression of X-linked genes was observed also in the gonadal primordia, which was comparable to that of the liver (Figs. 5a and 6c). Conversely, genes with significant sexual bias in their expression were enriched with X-linked genes among all the genes based on our gene model (38 X-linked genes out of 66 differentially expressed genes with false discovery rate < 0.01 (DEGs); odds ratio = 207.9, 95% CI 129.9 - 341.6).

These trends were also confirmed in the analysis using transcriptome assembly as a reference, which potentially incorporated genes excluded from the genome assembly (51 X-linked genes out of 85 DEGs; odds ratio = 155.2, 95% CI 98.0 - 249.1) (Extended Data Fig. 9b). Although a Y-linked gene *Zbtb39Y* exhibited the most pronounced male-biased expression (Fig. 6c), *Zbtb39X* and *-Y* are almost identical in their amino acid sequences (99.4%). Overall expression of *Zbtb39X/Y* was comparable between sexes, indicating their irrelevance to sex determination (Extended Data Fig. 9c). The majority of 38 X-linked genes included in DEGs were expected to be unrelated to sexual differentiation of gonadal primordia, according to their functional annotations (Fig. 6c and Extended Data Fig. 9b). Therefore, we again focused on the twelve X-linked genes implicated in intercellular signaling pathways (Extended Data Fig. 9b). Among them, *Acvr1b* and *Igfbp6* were expressed in a significantly female-biased manner. *Amhr2* exhibited no significant sexual difference in its expression, although it was expressed at a certain level (Fig. 6c, d, and Extended Data Fig. 9d).

We also explored genes that are well-characterized for their sex-specific role in gonads across bony vertebrates (*Amh*, *Amhr2*, *Cyp17a1*, *Dmrt1*, *Sox9*, *Foxl2*, and *Cyp19a1*)^60–62^ using RNA-seq data from juvenile gonads in addition to gonadal primordia. *Sox9*, *Foxl2*, and *Cyp19a1* were expressed in gonadal primordia, while *Dmrt1* and *Amh* were not detected (Fig. 6d). Unexpectedly, *Foxl2* and *Cyp19a1* exhibited comparable expression levels between sexes (Fig. 6d). In contrast, these genes were confirmed to be expressed in an expected direction in juvenile gonads; *Foxl2* and *Cyp19a1* were expressed with the female bias, while *Amh*, *Amhr2*, *Cyp17a1*, *Dmrt1*, and *Sox9* were expressed with the male bias (Fig. 6e).

## Discussion

Sex chromosomes undergo their lifecycle in evolution along the acquisition of a sex determining gene, degeneration, dosage compensation and turnover^10–13^. Recent studies have discovered the exceptions to this model^14,15,18–20^, which implies the previously unrecognized but critical aspects of sex chromosome evolution. In this study, we aimed to elucidate such aspects by investigating sex chromosome evolution of cartilaginous fishes.

Our analysis validated sex chromosome identification in multiple species spanning sharks and rays (Fig. 3a). Synteny analysis revealed the homology among the X chromosomes identified in individual species (Fig. 3b). This result indicated that the X chromosomes and their sex determining function originated before the common ancestor of sharks and rays. Based on the evolutionary timing of the shark-ray split^63^, the origin of these sex chromosomes can be traced back to around 300 mya, at least.

One recent study on the silver chimaera (*Chimaera phantasma*) genome reported its small X chromosome-like region, which showed its homology to autosomes in the epaulette shark and the zebra shark^64^. Hence, the origin can be traced back to the elasmobranch-chimaera split around 400 mya, at the oldest. This record is much older than the origins of the sex chromosomes of mammals, birds, anole lizards, and sturgeons that were previously known as the oldest to date among vertebrates (100 to 200 mya)^3,4,19,65^. This ancient origin is concordant with the extensive differentiation of sex chromosomes of sharks and rays, featuring short PARs as well as the scarce gene repertoires on the shark Y chromosomes (Figs. 3c and 4). One well-defined feature of heteromorphic sex chromosomes, such as mammalian X and avian Z chromosomes, is evolutionary strata, which are distinct genomic regions that reflect progressive loss of homologous recombination at different timings^66–68^. This was originally defined by stepwise changes in synonymous substitution rates between X-Y gametologs along their chromosomal positions^66^, and was also reflected by density of transposable elements and GC-content^68,69^, and potentially by heterozygosity because of the reduced population size. Owing to the massive loss of ancestral Y-linked genes, only clue for this structure is sequence properties of X chromosomes. Considering uniform, non-stepwise distributions of the heterozygosity and GC-content along the X chromosomes (Extended Data Fig. 3), evolutionary strata were not determined on the elasmobranch sex chromosomes. Even though the X1-added region of the red stingray is estimated to form a younger stratum compared with the rest of the X1 chromosome, they did not exhibit such differences. This implies that evolutionary strata were attenuated by the long evolutionary time after the latest recombination supression. Sex chromosomes in sharks and rays are represented by over 300 million years of antiquity with its stability once their differentiation has reached a certain degree.

Another remark of their sex chromosomes is incomplete dosage compensation. Although this was suggested in the white-spotted bamboo shark^27^, our choice of species allowed us to uncover its conservation from sharks to a ray (Fig. 5), consolidating the absence of complete dosage compensation widely in elasmobranchs. Early studies in model organisms once suggested that chromosome-level dosage compensation was a consequence of sex chromosome differentiation. Consistently, most of differentiated XY chromosomes studied thus far show chromosome-level dosage compensation (e.g. placental mammals, fruit flies, nematodes, and the green anole)^5,65,70,71^, while most of differentiated ZW chromosomes show incomplete dosage compensation (e.g. birds, snakes, and a trematode *Schistosoma mansoni*)^14,15,72–74^, implicating heterogamety in the evolution of dosage compensation^16,17^. Our finding in differentiated XY chromosomes provides robust counterexample to this dogma, prompting us to reconsider the model for dosage compensation.

Molecular basis of sex determination in cartilaginous fishes remained unexplored, which was tackled in the present study by utilizing rich sources of bamboo shark embryonic materials. Our histological analysis characterized the process of the sexual differentiation of gonadal primordia in cartilaginous fishes, which were synchronized with that of reproductive tracts and external genitalia (Fig. 6a, b). Transcriptomic analyses of embryonic gonads revealed that genes on the YDR were rarely expressed, while considerable numbers of genes on the XDR showed female-biased expression, because of the incomplete dosage compensation (Fig. 6c). Regarding gene repertoires, none of the orthologs of previously documented sex determining genes were located on sex chromosomes, except for *Amhr2* (Fig. 3d). *Amhr2* was located on PARs or autosomes and showed sexually comparable expression in embryonic gonads, suggesting its irrelevance to sex determination. These results indicate that gonadal sex determination in sharks and rays is accomplished by the differential expression from halved dosages of X-linked genes that have never been documented as master sex determining genes. Whether a single gene or multiple genes trigger the sex determination cascade, *Acvr1b* and *Igfbp6* are promising candidates for the master sex determining gene because of their significantly female-biased expression with their potential function in gonadal development (Fig. 6c, d). ACVR1B (also known as ALK- 4) is a receptor for multiple TGF-β ligands including activins^75^. Activins are involved in a wide range of developmental processes^75,76^ including ovarian follicle formation^77–79^, testicular development^45–48^, germ cell proliferation^46,80^, germ cell differentiation^47,81^, and gonadal steroidogenesis^82,83^. IGFBP6 is a member of IGF binding proteins, which modulate signal transduction activity by IGF ligands^84,85^. IGF signaling pathway is also implicated in a wide range of developmental processes including ovarian development^86^, testicular development^53^ and germ cell differentiation^54,87^. Considering their pleiotropic roles especially in gonadal development, we hypothesize that either or both of X-linked *Acvr1b* and *Igfbp6* canalize(s) the ovarian morphogenesis of bipotential gonadal primordia. Sex determination in sharks and rays may be underpinned by gene-dosage-dependent mechanism with unique molecules distinct from other vertebrate lineages.

In contrast, molecules involved in the maintainance of gonadal sex identity (*Foxl2*, *Cyp19a1*, *Amh*, *Amhr2*, *Cyp17a1*, *Dmrt1*, and *Sox9*)^60–62^ were expressed in the similar pattern between cartilaginous fishes and bony vertebrates. Our transcriptomic analysis of juvenile bamboo shark gonads revealed that these genes were expressed with the sexual biases, while most of these genes lacked such biases at the timing of sex determination (Fig. 6d, e). These observations further highlight the uniqueness of the sex determination mechanism in sharks and rays, which might be composed of two temporally separated phases: the upstream cascade composed of novel factors and the downstream network operated by well-conserved factors. This unique example will expand our view on the evolutionary plasticity of sex determination mechanism; how the diverse molecules can be linked to sexually antagonistic network for vertebrate sex determination.

In conclusion, our study discovered the oldest vertebrate sex chromosome without global dosage compensation. Our results delineate the unique molecular mechanism of sex determination in sharks and rays distinct from other vertebrate lineages. These findings advance our understanding on the sex chromosome evolution and shed light on the evolutionary plasticity of vertebrate sex determination.

## Methods

### Animal sampling

Samples of the bamboo shark, cloudy catshark, red stingray and elephant fish were provided by the public aquariums of Osaka Aquarium Kaiyukan, Enoshima Aquarium and Maxell Aqua Park Shinagawa, Shimoda Aquarium and Aquarium Facility of RIKEN Center for Developmental Biology, Atmosphere and Ocean Research Institute (AORI) of University of Tokyo, or purchased from commercial pet supplier (see supplementary table). Animals were sacrificed after anesthetizing by cold sea water or MS-222. For bamboo shark embryos, fertilized eggs provided by the public aquariums were cleaned by peeling off opaque surfaces and maintained in aquarium tanks at 25 ℃ until they reached the optional developmental stage. Staging was based on the table for embryonic development^58^. Animals were sexed by the presence and absence of the claspers, male-specific structures on pelvic fins. The sex of bamboo shark embryos before clasper formation was determined by PCR targeting the 400 bp genomic region on the YDR, for which tissue lysates were prepared by boiling tissue fragments in 50mM NaOH at 95 °C, followed by 1M Tris HCl (pH8.0) addition. The supernatant was used for the PCR with KOD FX Neo polymerase (Takara) and the primer pair of 5′- cctaagagcatcaccttacc-3′ and 5′-gagtactcatagctgcaagg-3′. Animal handling and sample collections at the aquariums were conducted by veterinary staff without restraining the individuals, in accordance with the Husbandry Guidelines approved by the Ethics and Welfare Committee of Japanese Association of Zoos and Aquariums. All other experiments were conducted in accordance with the Guideline of the Institutional Animal Care and Use Committee (IACUC) of RIKEN Kobe Branch (Approval ID: H16-11), National Institute of Genetics (Approval ID: R5-14 and R6-13), or the Guideline for Care and Use of Animals at the University of Tokyo.

### Genome sequencing

High molecular weight DNA was extracted from the liver of a 20 cm-long male juvenile brownbanded bamboo shark (sChiPun1) and the blood of an adult female cloudy catshark (sScyTor2) using a NucleoBond AXG column (Macherey-Nagel), which was followed by purification with phenol-chloroform. The concentration of the extracted DNA was measured by Qubit 4 with the Qubit dsDNA HS Assay Kit (ThermoFisher), and their size distribution was analyzed by TapeStation 2100 with the Genomic DNA Screen Tape (Agilent Technologies) to ensure high integrity. A SMRT sequence library was constructed with an SMRTbell Express Template Prep Kit 2.0 (Pacific Biosciences) and was sequenced in a single 8M SMRT cell on a PacBio Sequel IIe system (Pacific Biosciences) at Kazusa DNA Research Institute for bamboo shark and on a PacBio Sequel II system at National Institute of Genetics for cloudy catshark. The sequencing output was processed into a total of 160 Gb and 230 Gb HiFi reads for the bamboo shark and cloudy catshark, respectively. Genomic DNA for whole genome resequencing was extracted from the livers, spleens and muscles of the bamboo shark, cloudy catshark, red stingray and elephant fish using Monarch Genomic DNA Purification Kit (New England Biolabs), NucleoBond AXG column (Macherey-Nagel), Qiagen DNeasy Blood & Tissue Kit (QIAGEN), and the phenol/chloroform method previously described^31^ (see supplementary table). Library preparation and sequencing were performed by Novogene Co., Ltd. via NovaSeq 6000 platform (Illumina) at PCR-free paired-end 150 bp strategy (PE150).

### Hi-C

The Hi-C library was prepared using the liver (bamboo shark) and muscle (cloudy catshark) of the individual used for HiFi read production, according to the iconHi-C protocol^88,89^ employing restriction enzymes DpnII for the bamboo shark and DpnII and HinfI for the cloudy catshark. The prepared libraries were sequenced on a HiSeq X sequencing platform (Illumina) at PE150. Hi-C reads were trimmed by fastp v0.23.4^90^ and mapped onto contigs using HiC-Pro v3.1.0^91^ with the MAPQ threshold 1.

### Genome assembly

The residual adapter sequences in the HiFi genomic reads were removed by HiFiAdapterFilt^92^. Processed HiFi reads were assembled into contigs using Hifiasm v0.16.1^93^. These contigs were scaffolded by YaHS v1.2^94^ with the option ‘--no-contig-ec --no-scaffold-ec -q 1 -e GATC --no-mem-check -r 2000,5000,10000,20000,50000,100000,200000,500000,1000000,2000000,5000000,100 00000,20000000,50000000,100000000,200000000,500000000’ for the bamboo shark and ‘-q 1 -e GATC,GANTC --no-mem-check -r 2000,5000,10000,20000,50000,100000,200000,500000,1000000,2000000,5000000,100 00000,20000000,50000000,100000000,200000000,500000000’ for the cloudy catshark.

The resulting contact maps were processed with the juicer pre command in YaHS and juicer_tools.1.9.9, and manually curated using Juicebox v1.11.08^95^ and converted into final genome assemblies via YaHS juicer post command or normalized and visualized by HiCExplorer v3.7.2^96^. Mitochondrial genome sequence of the bamboo shark was assembled from HiFi genomic reads using MitoHiFi v3.0.0^97^. The resulting assemblies were evaluated for completeness and continuity by gVolante v2.0.0^98^ using BUSCO v5^99^ with OrthoDB v10 Vertebrata peptide sequence dataset^100^.

### Repeat and gene annotation

Species-specific repeat libraries were constructed by RepeatModeler v2.0.5^101^ using the genome assemblies of the bamboo shark and cloudy catshark genomes. Repetitive sequences in the genome assemblies were detected by RepeatMasker v4.1.6 using the custom repeat libraries obtained above and soft-masked into lower cases with options ‘- nolow -xsmall’. Gene models on the bamboo shark, cloudy catshark, and shortfin mako genomes were constructed by the Braker v3.0.6^102^ pipeline, whose Augustus v3.4.0 was replaced with v3.5.0. Publicly available transcriptome data^31,103^ (see supplement) for the three species, our original transcriptome data for the bamboo shark and OrthoDB v10 Vertebrata peptide sequence dataset were used as the hints for gene model construction. Functional annotation for coding sequences was performed by EnTAP v1.0.1^104^ using RefSeq vertebrate_mammalian, RefSeq other_vertebrate, RefSeq invertebrate, UniProt Swiss-Prot and UniProt TrEMBL peptide sequence databases as the reference (downloaded on 24 January 2024). Distribution of interspersed repeats were calculated at non-overlapping 10 kb genomic windows using bedtools v2.31.0^105^ ‘makewindows’, ‘coverage’ subcommands and the RepeatMasker output. Tandem repeats on the bamboo shark Y chromosome were detected and classified based on the unit sequence and repeating number using TRASH v1.2^106^.

### X chromosome identification and characterization

The reference genome and its gene model of the red stingray were adopted from our parallelly conducted project (under NCBI BioProject PRJNA1206076). Short genomic reads from our experiments and public databases^33,35,36^ (thorny skate dataset from GenomeArk repository with sex identity shared personally by the VGP team) were mapped onto reference genomes using bwa-mem2 v2.2.1^107^ (see data sources in supplementary tables). Read depth was counted for 10-kb non-overlapping genomic windows using mosdepth v0.3.3^108^ and normalized by the average value of these windows. Heterozygous site was detected by bcftools v1.9^109^ mpileup/call using FORMAT/GT tag and its frequency was calculated at 100-kb non-overlapping genomic windows. For both analyses, each individual was treated independently. GC-content of the X chromosomes was calculated for 10 kb non-overlapping genomic windows using bedtools v2.31.0^105^ ‘makewindows’ and ‘nuc’ subcommands.

### Synteny comparison

Synteny blocks on X chromosomes were calculated and visualized by GENESPACE v1.3.1^110^ using positions and peptide sequences of protein-coding genes. Our original gene models were used for the bamboo shark, cloudy catshark, red stingray and shortfin mako and RefSeq gene models were adopted for the other species. For gene-by-gene synteny visualization, orthology information was extracted from intermediate files from GENESPACE and visualized by NGenomeSyn v1.4.1^111^. Homology between X and Y chromosomes, or Y chromosomes between two species were detected and visualized by ModDotPlot v0.8.7^112^ in static mode.

### Ortholog search

Cartilaginous fish orthologs of *Dmrt1*, *Foxl2*, *Sox3*, *Dhh*, *Gdf11*, *Wnt1*, *Wnt6b*, and *Wnt10b* genes were identified by pangene function in GENESPACE v1.3.1 using non-cartilaginous fish vertebrate datasets (human, chicken, medaka, northern pike, fugu, and turquoise killifish) in addition to our study species. For the detailed analysis of ten genes (*Amh*, *Amhr2*, *Gdf6*, *Gsdf*, *Acvr1b*, *Acvr1l*, *Gli1*, *Igfbp6*, *Stat2*, and *Stat6*), molecular phylogenies were inferred using MAFFT v7.525^113^ with ‘--auto’, TrimAl v1.4.rev15^114^ with individualized trimming settings and IQ-TREE v2.2.6^115^ with ‘-m MFP -B 10000’ for alignment, trimming and tree inference, respectively^116,117^.

Thresholds for gap trimming were set ‘-gt’ as 0.7, 0.1, 0.2, 0.1, 0.1, and 0.1 for *Amhr2*, *Amh-Gsdf*, *Gdf6*, *Acvr1b-Acvr1l*, *Gli1* and *Stat2-Stat6*, respectively, and automatically set ‘-automated1’ for *Igfbp6*. Peptide sequences were collected by NCBI BLASTP search for the NCBI RefSeq database, manual collection from Ensembl, BLASTP (BLAST 2.14.0+) search for our original gene models and TBLASTN search in the Squalomix sequence archive^31,118^. In addition, several peptide sequences were collected by peptide to nucleotide similarity search against genomic sequences using miniprot v0.12-r237^119^ with the peptide sequences of most closely related species (sequences uploaded to figshare).

### Y chromosome microdissection followed by sequencing

Metaphase chromosome spreads were prepared from the primary culture of fibroblasts obtained from a stage 29 male bamboo shark embryo as described previously^23^. For microdissection, we used a phase-contrast microscope Zeiss Axiovert.A1 (Zeiss) equipped with Eppendorf InjectMan NI 2 micromanipulator (Eppendorf). Glass needles were made from 1.0 mm diameter capillary glass using a glass capillary puller, P-10 Micropipette Puller (Narishige) and sterilized by ultraviolet irradiation. Individual Y chromosomes in each metaphase spread were scratched with the glass needles and transferred into separate 0.2-ml PCR tubes. Y chromosome DNAs were amplified using the Illustra Single Cell GenomiPhi DNA Amplification kit (Cytiva). The accuracy of chromosome dissection was evaluated by fluorescent *in situ* hybridization according to our previous protocol^23^. Briefly, the Y chromosomal amplicons were labeled with biotin 16-dUTP using a nick translation kit (Roche Diagnostics) and hybridized with metaphase spreads. These spreads were treated by avidin conjugated with Alexa Fluor 488 (Thermo Fisher Scientific-Molecular Probes) and mounted with the Vectashield mounting medium with DAPI (Vector Laboratories). The amplicons were sheared using Covaris E220 with the condition targeting 200 bp in 50 μL sample volume in microTUBE (COVARIS), and subjected to size selection with AMpure XP (Beckman Coulter) to retrieve DNA larger than 100 bp. Sequencing libraries were prepared from three micro-dissected samples, with KAPA LTP Library Preparation Kit (Roche Diagnostics) and sequenced by HiSeq 1500 platform (Illumina) at PE251. Sequencing reads were mapped onto the bamboo shark reference genome by bwa-mem2, and read counts were calculated by mosdepth per scaffold basis. The identified scaffold was concatenated with the PAR sequence in the bamboo shark X chromosome and used for the following analyses as the Y chromosome.

### Virtual 4C analyses

For the bamboo shark YDR, bamboo shark Hi-C reads were mapped onto the reference genome using HiC-Pro v3.1.0^91^ without any MAPQ thresholds, and valid read pairs in the HiC-Pro intermediate file were filtered by the mapping position and MAPQ; either of read pairs were assigned on the YDR and the other read in the pair had MAPQ > 1. Filtered reads were counted at non-overlapping 100 kb genomic windows. For the bamboo shark *Amhr2* locus, Hi-C reads were mapped onto the modified reference genome including a 14-kb HiFi genomic read containing a part of *Amhr2* CDS, which was obtained using minimap2 v2.26-r1175^120^ with default parameters and *Amhr2* CDS derived from the previously published transcriptome assembly. Valid read pairs were filtered only by the mapping position; either of read pairs were assigned on the *Amhr2*- containing HiFi read. Filtered reads were counted per chromosome basis.

### Histological analyses

Bamboo shark embryos at stages 30, 31, and 32 were fixed in Bouin’s fixative at room temperature overnight, dehydrated by 70% ethanol, 100% ethanol, xylene, and embedded in paraffin blocks. Paraffin blocks were sectioned at 7 μm. Paraffin sections were washed with xylene, ethanol and stained with hematoxylin and eosin. Slides were enclosed by MGK-S (Matsunami) and observed by Primostar 3 (Zeiss).

### Transcriptome analyses

Total RNA was extracted from livers of the bamboo shark, cloudy catshark, and red stingray as well as bamboo shark developing gonads from stage 31 embryos and juvenile individuals using the Direct-zol RNA extraction kit (Zymo Research), TRIzol (ThermoFisher Scientific) or ISOGEN (Nippongene), followed by DNaseI treatment (ThermoFisher Scientific). RNA quality was evaluated by Bioanalyzer 2100 with the RNA 6000 nano kit or Tapestation D2200 with the High sensitivity RNA ScreenTape (Agilent Technologies). Total RNA from bamboo shark livers and gonads were processed with the TruSeq Stranded mRNA Prep kit (Illumina) and sequenced with HiSeq 1500 platform at PE127 or HiSeq X platform (Illumina) at PE150 through Azenta Life Sciences. Library preparation and sequencing from the total RNA of red stingray livers and cloudy catshark livers were performed by Novogene Co., Ltd. via NovaSeq 6000 platform (Illumina) at PE150. Sequencing reads were trimmed by fastp v0.23.4^90^, mapped onto reference genomes using hisat2 v2.2.1^121^ and counted by featureCounts v2.0.6^122^ using our original gene models. For male-female expression ratio, read counts were normalized by total mapped read counts into CPM, and the ratio was calculated from the CPM averaged by each sex. The statistical analysis for DEGs was performed via edgeR v3.42.4^123^. Transcriptome assembly-based expression analysis was performed via Trinity v2.15.1^124^ utilizing trimmomatic^125^ module for read trimming and align_and_estimate_abundance.pl for read counting^126^. Reading frame detection and functional annotation for transcriptome assemblies was performed by EnTAP v1.0.1^104^ with TransDecoder v5.7.1 using RefSeq vertebrate_mammalian, RefSeq other_vertebrate, RefSeq invertebrate, UniProt SwissProt and UniProt TrEMBL peptide sequence databases as the reference (downloaded on 24 January 2024). The genomic location of each transcript isoform was assigned by minimap2 v2.26-r1175^120^ with ‘-x splice:hq -N 1’ followed by best hit search among isoforms with MAPQ, query cover and match length. Gene-based count matrix was used for DEG analysis in the same method as the genome assembly-based approach.

## Data availability

Sequence data and genome assemblies are deposited in NCBI under the accession number GCA_047496795.1 (bamboo shark) and GCA_047496885.1 (cloudy catshark) in the BioProject: PRJNA1202043. The other data associated with this study are available at https://figshare.com/projects/sharksexchr/232328.

## Acknowledgments

The authors thank Eric Jarvis, Olivier Fedrigo and Gavin J. P. Naylor for sharing metadata of phenotypic sex of thorny skate samples and Susumu Hyodo for sample provision of cloudy catshark. Our gratitude extends to Osamu Nishimura, Kaori Tatsumi, Chiharu Tanegashima for technical assistance, Kazumi Matsubara and Yoshiyuki Seki for assistance in chromosome dissection, and Kazuaki Yamaguchi, Yasuhisa Kobayashi, Szu-Hsuan Lee, Yann Guiguen, Manfred Schartl, Luohao Xu, Kateryna Makova for insightful discussion. This work was supported by Junior Research Associate (JRA) of RIKEN to Y.O. and funded by intramural budgets granted by RIKEN and the National Institute of Genetics, as well as JSPS KAKENHI Grants Number 20H03269 and 16H06279 (PAGS) to S.K., 21K06286 to Y.U., and 23KJ1002 (Grant-in-Aid for JSPS Fellows) to T.N.. Computations were partially performed on the NIG supercomputer at ROIS National Institute of Genetics.

---

**Extended Data Fig. 1.**
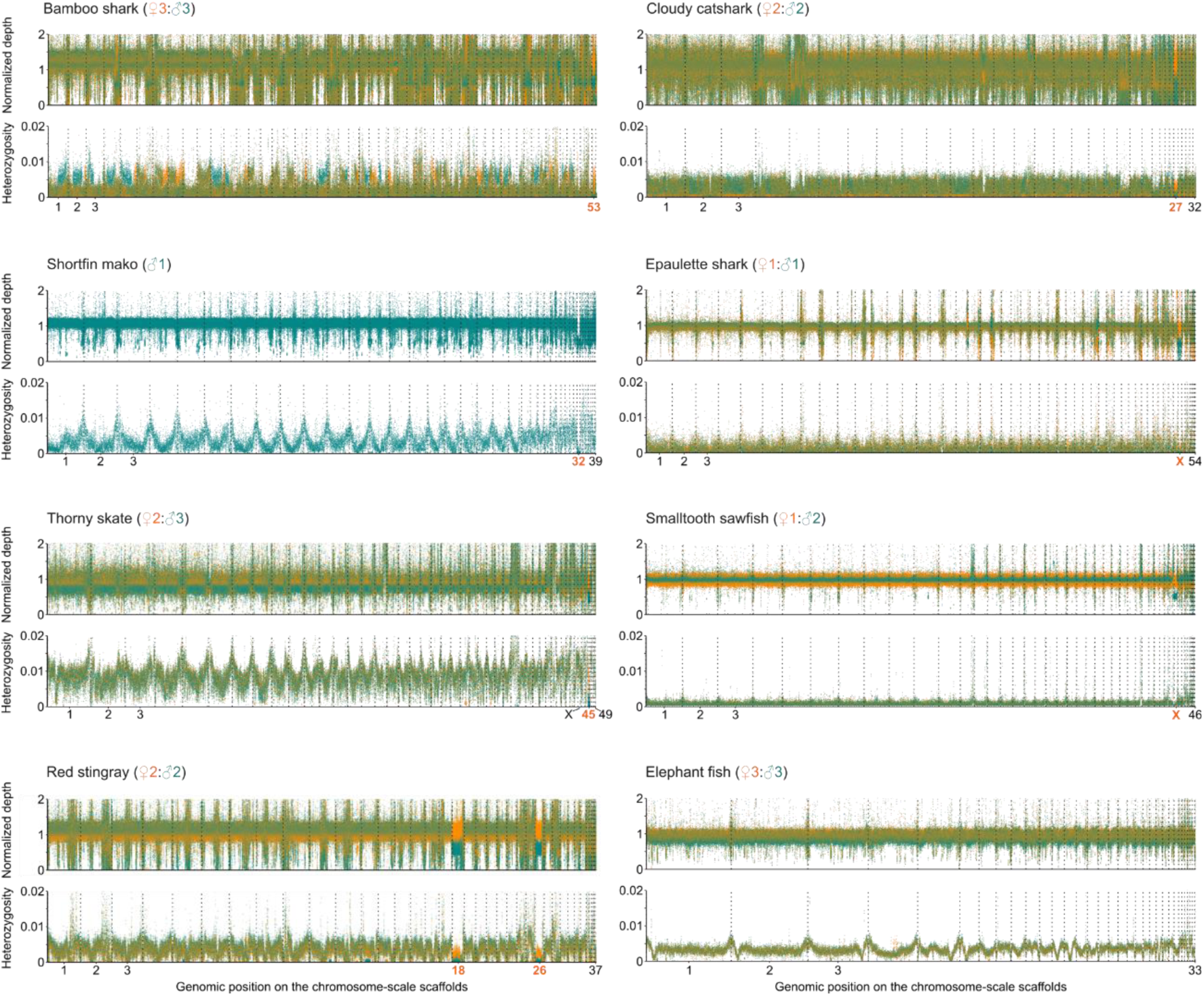
Male-female comparison of genome sequence features. Normalized sequencing depth and heterozygosity of chromosome-scale scaffolds were derived from whole genome resequencing of male and female individuals. These metrics were calculated for non-overlapping windows of 10 kb and 100 kb, respectively. Sequencing depth was normalized by its whole genomic average. Scaffolds were ordered by length.

**Extended Data Fig. 2.**
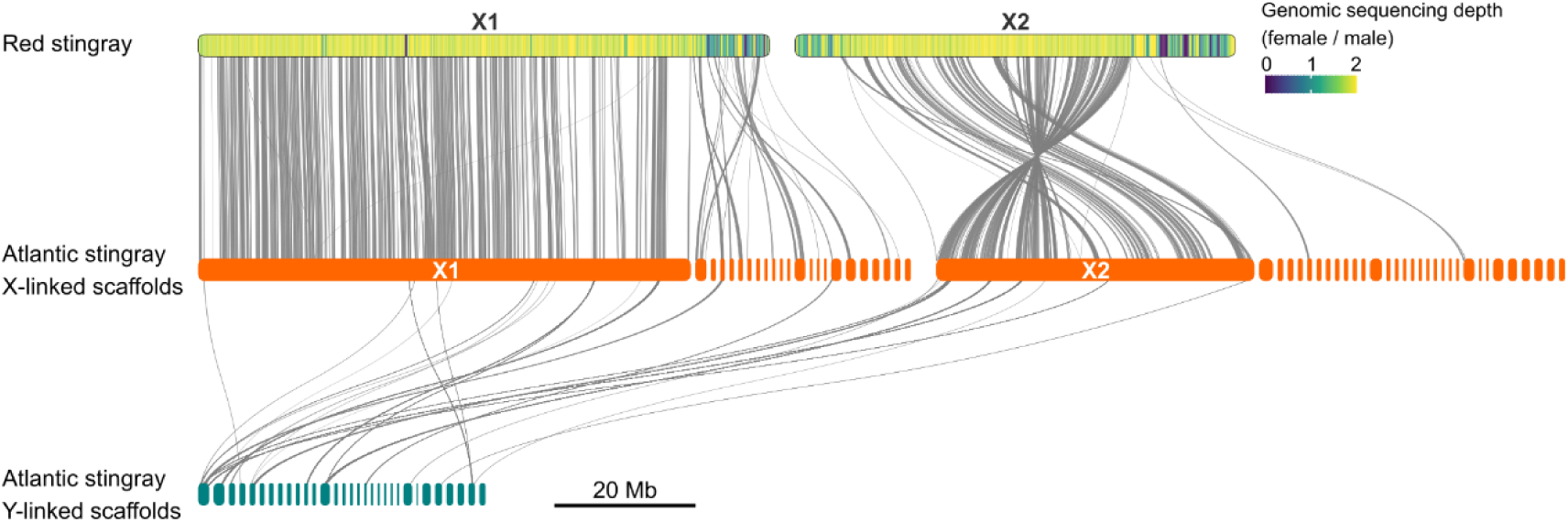
Gene repertoires shared in common between X1/X2 and Y chromosomes in the two myliobatiform species. Orthology, as well as X-Y correspondence for the Atlantic stingray, are visualized as gray lines. Atlantic stingray scaffolds designated as X1, X2, and Y chromosomes are shown as clusters of orange and blue bars. Female-to-male ratio of genome sequencing depth for 100-kb non-overlapping windows are shown with variable colors on the X1 and X2 chromosomes in the red stingray.

**Extended Data Fig. 3.**
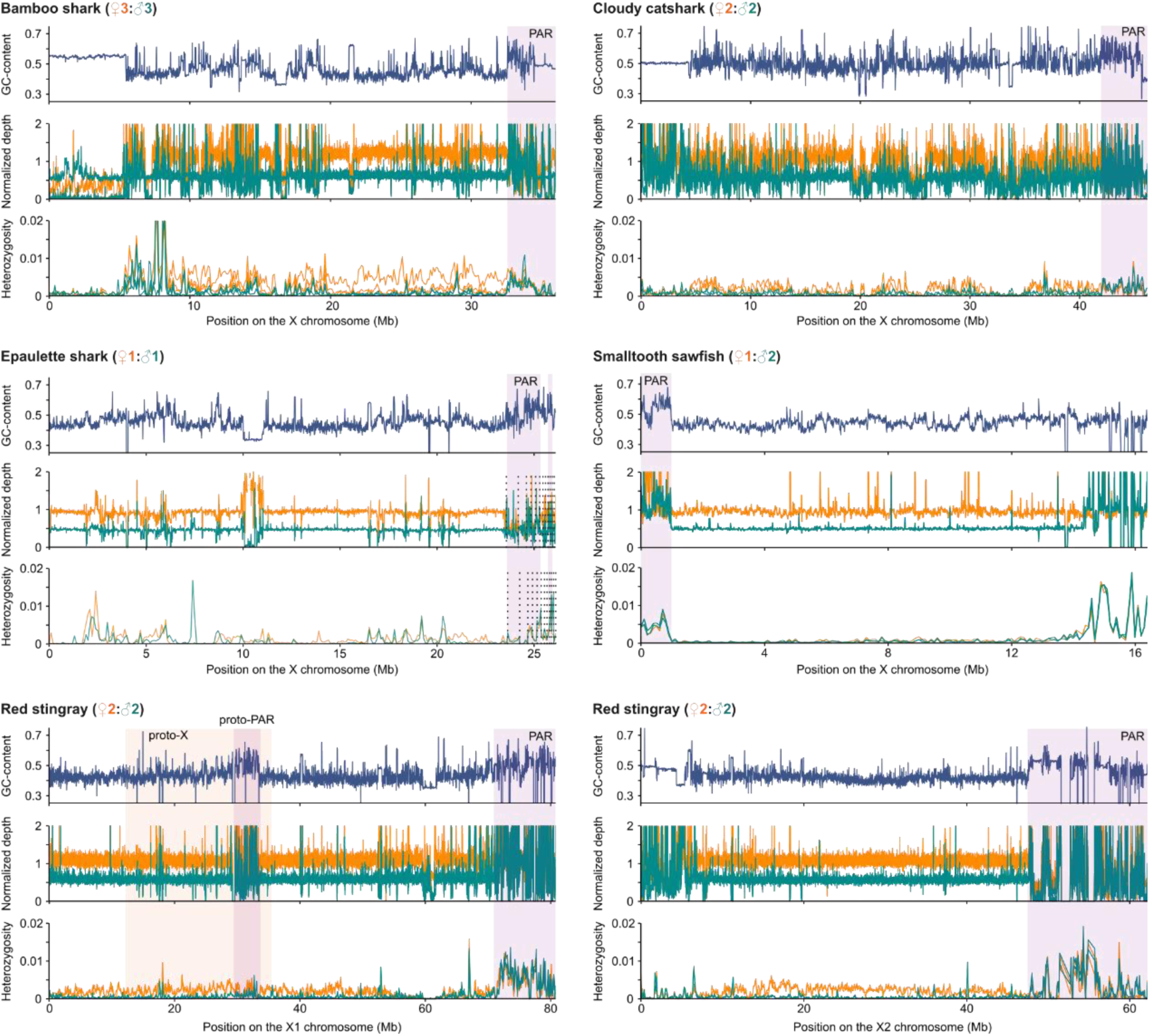
Sequence properties of X chromosomes. GC-content, normalized sequencing depth and heterozygosity are shown at non-overlapping windows of 10 kb, 10 kb, 100 kb, respectively. For the epaulette shark, multiple small scaffolds registered as parts of the X chromosome are concatenated. Purple shades indicate PARs identified with GC-content and normalized sequencing depth.

**Extended Data Fig. 4.**
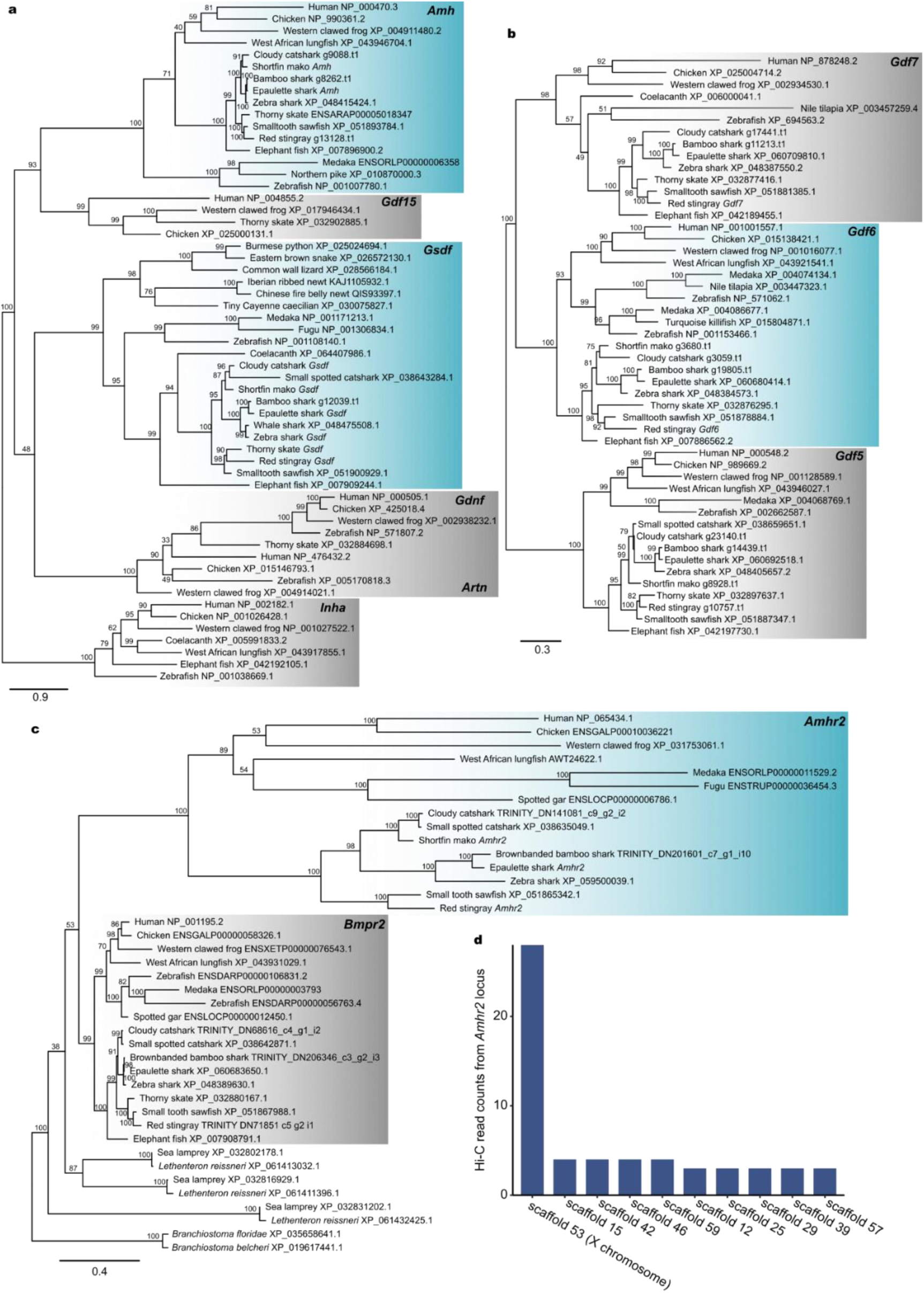
Orthology assessment of the typical master sex determining genes. **a-c**, Phylogenetic trees of vertebrate *Amh* and *Gsdf* (**a**), *Gdf6* (**b**), and *Amhr2* (**c**) inferred by the maximum likelihood method. The values at nodes indicate bootstrap probabilities with 10,000 resamplings. **d**, Total read counts of each scaffold at the virtual 4C analysis using the bamboo shark *Amhr2* as the target sequence. Scaffolds were ordered by amounts of read counts.

**Extended Data Fig. 5.**
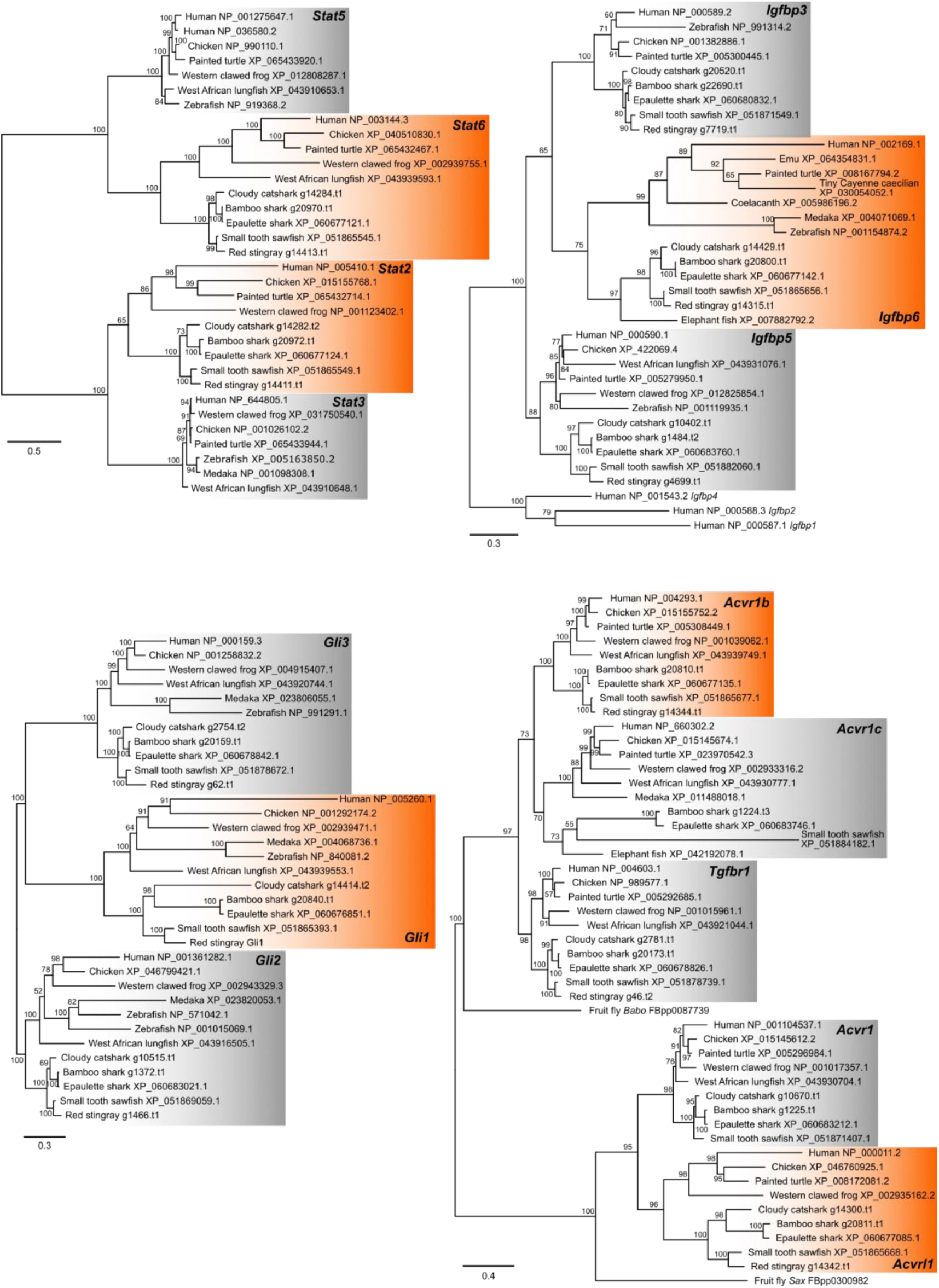
Molecular phylogeny of X-linked genes related to intercellular signaling pathways. Phylogenetic trees of vertebrate *Stat2, -6* (**a**), *Gli1* (**b**), *Igfbp6* (**c**) and *Acvr1b* and *Acvrl1* (**d**) were inferred by the maximum likelihood method, respectively. The values at nodes indicate bootstrap probabilities with 10,000 resamplings.

**Extended Data Fig. 6.**
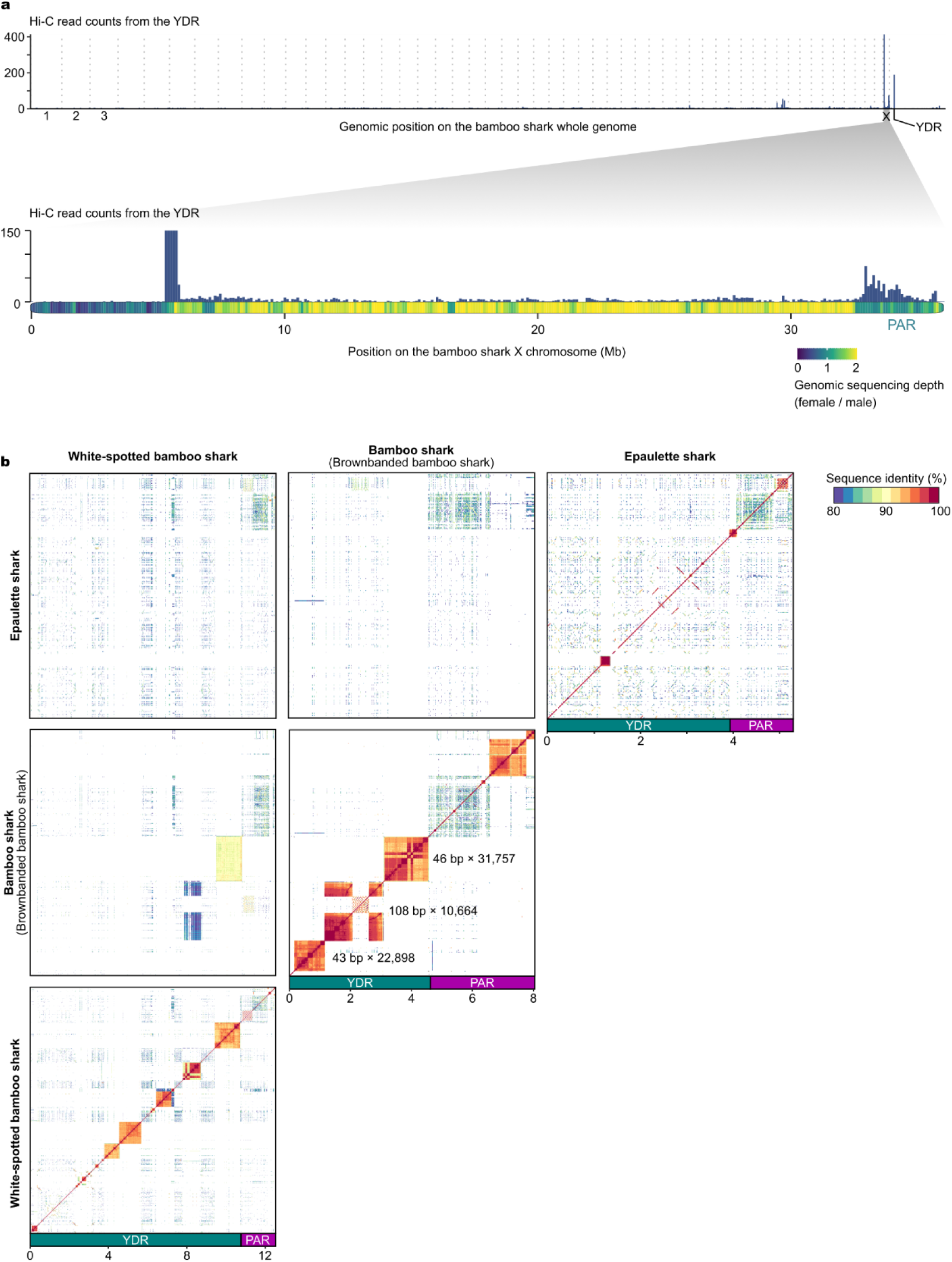
Identification of shark Y chromosomes and their degeneration. **a**, Read counts for 100-kb non-overlapping windows at the virtual 4C analysis using the bamboo shark YDR as the target sequence. Scaffolds were ordered by length at the whole genome view. Female-to-male ratio of genome sequencing depth for 100-kb non-overlapping windows are shown with variable colors on the X chromosome. **b**, Self- and cross-comparison of sequence similarity among three shark Y chromosomes. Sequence similarities were computed at 10 kb non-overlapping windows. Numbers in the self-similarity plot of the bamboo shark Y chromosome represent unit lengths and repeating numbers of tandem repeats on the YDR.

**Extended Data Fig. 7.**
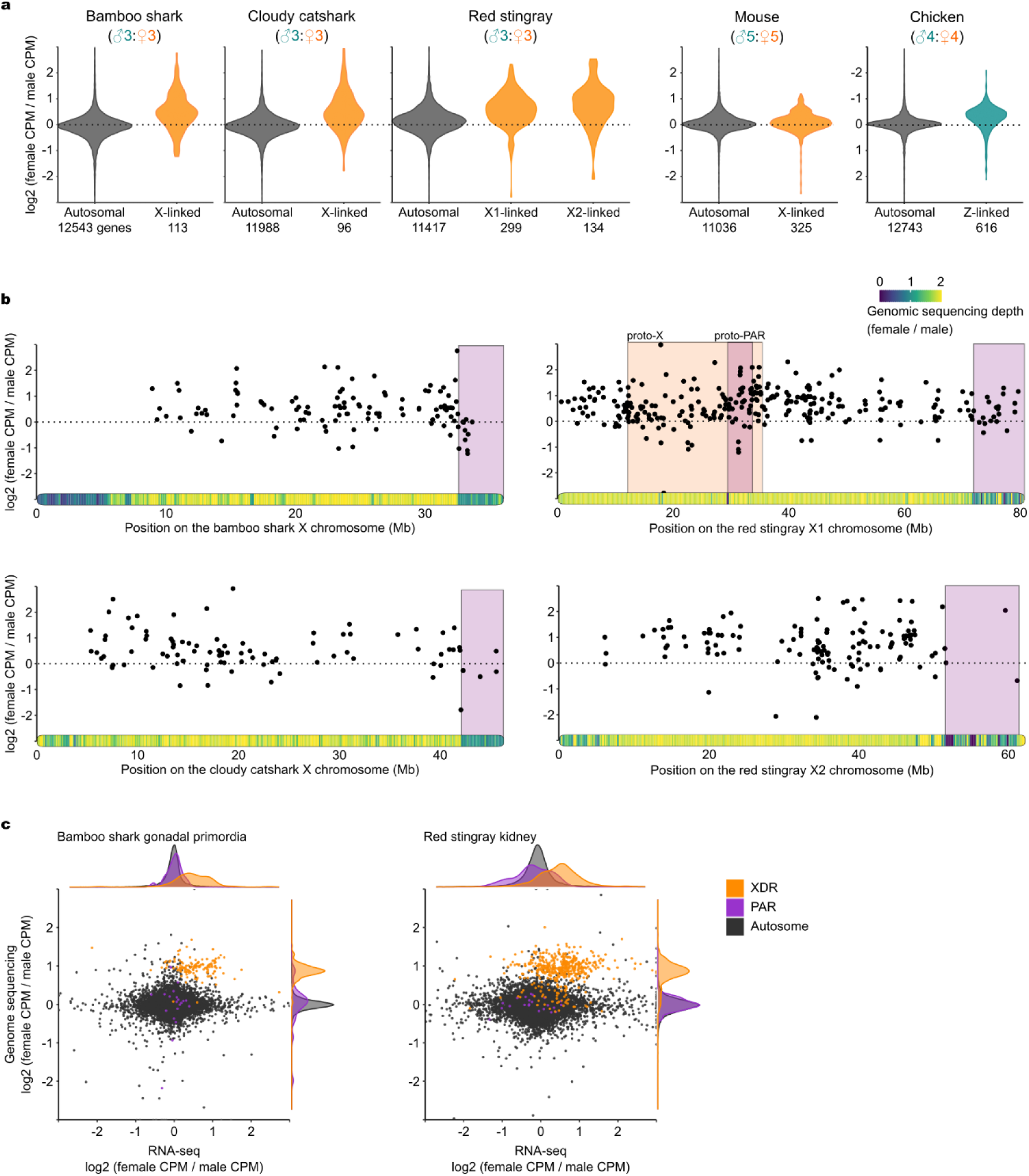
Dosage dependent expression of X-linked genes. **a**, Distributions of the female-to-male ratio of gene expressions in the liver. Numbers below each column represent numbers of the genes analyzed. RNA-seq data for mouse and chicken, included for comparisons, were produced previously^55,56^. Note that the vertical axis for the chicken with the ZW system is inverted **b**, Sexual biases in gene expression along the position on X chromosomes of the three species. Each dot represents each gene. All values are averaged for individuals of each sex for each gene. **c**, Sexual biases in gene dosage and gene expression in gonadal primordia of the bamboo shark and kidneys of the red stingray.

**Extended Data Fig. 8.**
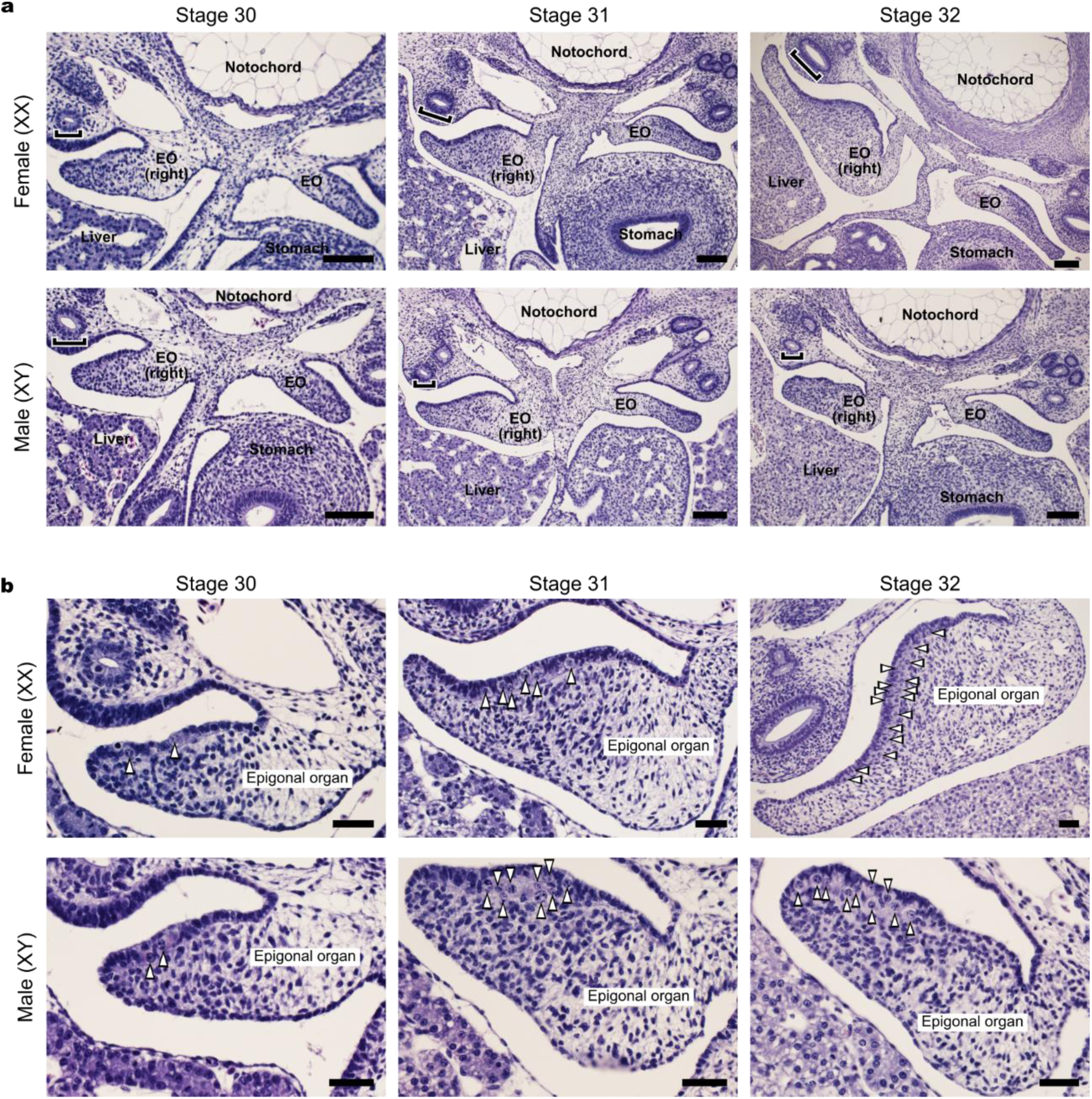
Sexual dimorphism in bamboo shark gonadal primordia. **a**. **b**, Transverse sections of bamboo shark trunks at the embryonic stage 30, 31, and 32. Images in **b** are enlarged from right gonadal primordia in **a**. Sections were stained with hematoxylin and eosin. EO: epigonal organ, brackets: Müllerian duct, arrowheads in b: germ cell. Scale bars, 100 μm (**a**), 40 μm (**b**).

**Extended Data Fig. 9.**
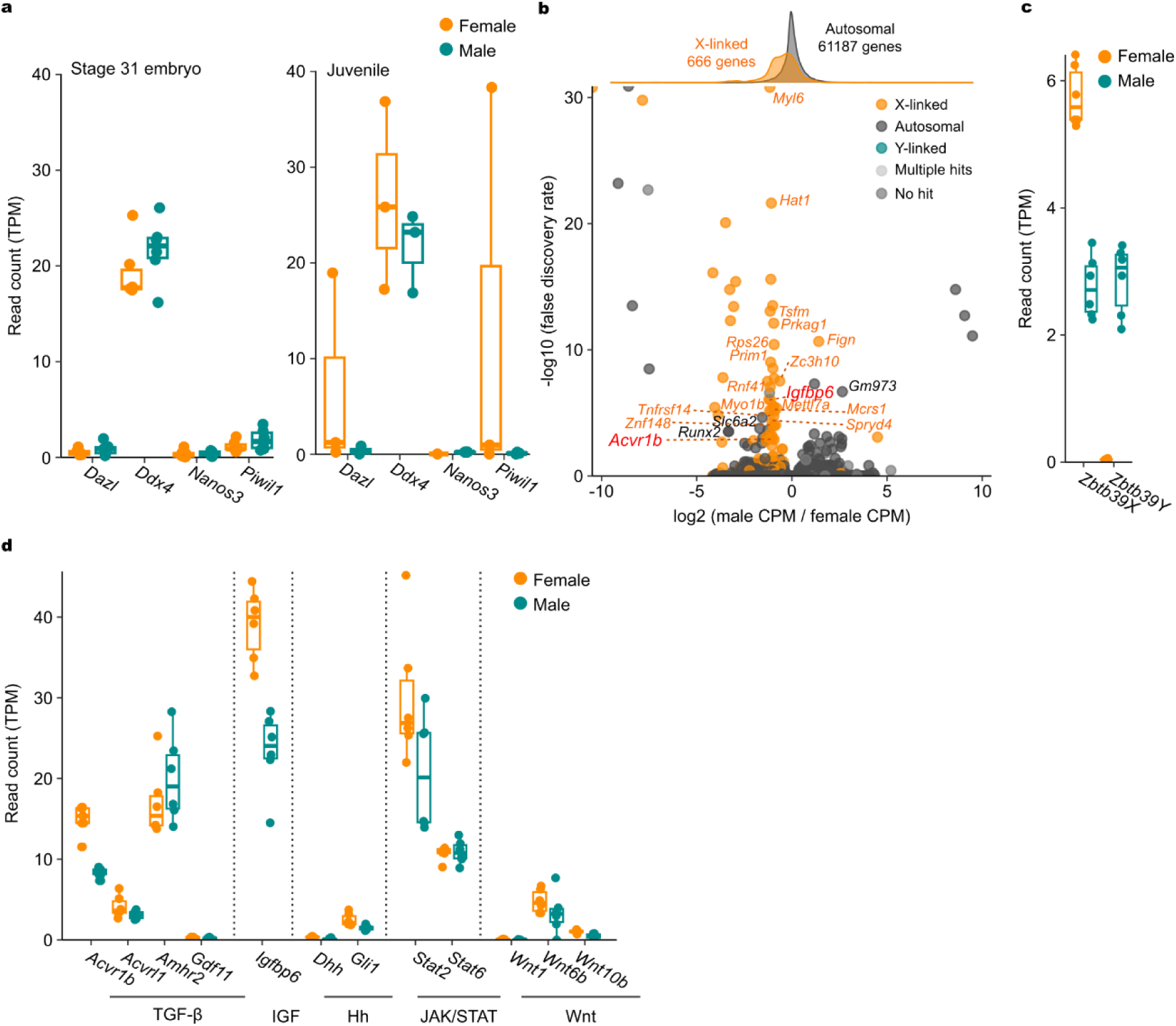
Quality of the bamboo shark transcriptomic data. **a**, Expression levels of genes implicated in germ cell functions in embryonic and juvenile samples. **b**, Differentially expressed genes between sexes in gonadal primordia. Transcriptomic reads are counted using transcriptome assembly as a reference. Gene annotations are based on similarity search of peptide sequences. See Methods for details. **c**, Expression levels of *Zbtb39* gametologs in embryonic samples. **d**, Expression levels of twelve X-linked genes implicated in intercellular signaling pathways in gonadal primordia. Boxplot elements are defined as follows: center line, median; box limits, upper and lower quartiles; whiskers, maximum and minimum within 1.5×IQR from hinges.

